# The comparative biogeography of Philippine geckos challenges predictions from a paradigm of climate-driven vicariant diversification across an island archipelago

**DOI:** 10.1101/395434

**Authors:** Jamie R. Oaks, Cameron D. Siler, Rafe M. Brown

## Abstract

A primary goal of biogeography is to understand how large-scale environmental processes, like climate change, affect diversification One often-invoked but seldom tested process is the “species-pump” model, in which repeated bouts of co-speciation are driven by oscillating climate-induced habitat connectivity cycles. For example, over the past three million years, the landscape of the Philippine Islands has repeatedly coalesced and fragmented due to sea-level changes associated with glacial cycles. This repeated climate-driven vicariance has been proposed as a model of speciation across evolutionary lineages codistributed throughout the islands. This model predicts speciation times that are temporally clustered around the times when interglacial rises in sea level fragmented the islands. To test this prediction, we collected comparative genomic data from 16 pairs of insular gecko populations. We analyze these data in a full-likelihood, Bayesian model-choice framework to test for shared divergence times among the pairs. Our results provide support against the species-pump model prediction in favor of an alternative interpretation, namely that each pair of gecko populations diverged independently. These results suggest the repeated bouts of climate-driven landscape fragmentation has not been an important mechanism of speciation for gekkonid lizards on the Philippine Islands.

## Introduction

Understanding how environmental changes affect diversification is an important goal in evolutionary biology, biogeography, and global change biology. Environmental processes that operate at or above the level of communities can simultaneously cause speciation or extinction across multiple evolutionary lineages, and thus have a pronounced effect on the diversity and distribution of species. Island archipelagos that harbor diverse communities of co-distributed lineages and have a relatively well-understood geological history present powerful systems for understanding such shared processes of diversification (Gillespie, 2007; Losos and Ricklefs, 2009; Vences et al., 2009; Brown et al., 2013). The Philippine archipelago represent such a model system, with more than 7,100 islands that arguably harbor the highest concentration of terrestrial biodiversity on Earth (Catibog-Sinha and Heaney, 2006; Brown and Diesmos, 2009; Heaney and Regalado, 1998; Brown et al., 2013); how, when, and by which mechanisms this diversity accumulated has piqued the interest of evolutionary biologists since the early development of the field of biogeography (Wallace, 1869; Huxley, 1868; Dickerson, 1928; Diamond and Gilpin, 1983; Brown, 2016; Lomolino et al., 2016).

The landscape of the Philippines has experienced a complex history. Climatological oscillations, primarily during the Pleistocene, led to the repeated formation and fragmentation of Pleistocene Aggregate Island Complexes (PAICs; Inger, 1954; Heaney, 1985; Brown and Diesmos, 2002, 2009; Esselstyn and Brown, 2009; Siler et al., 2010; Brown and Siler, 2014; Lomolino et al., 2016). During lower sea levels of glacial periods, islands coalesced into seven major landmasses (PAICs) that were fragmented into individual islands during interglacial periods. These climate-driven cycles have occurred at least six times during the last 500,000 years (Rohling et al., 1998; Siddall et al., 2003; Spratt and Lisiecki, 2016), with additional cycles occurring in the late Pliocene and early Pleistocene (Haq et al., 1987; Miller et al., 2005).

For nearly three decades, the repeated formation and fragmentation of PAICs has been a prominent model of diversification in the Philippines (Inger, 1954; Heaney, 1985; Brown and Guttman, 2002; Evans et al., 2003; Heaney et al., 2005; Roberts, 2006; Linkem et al., 2010a; Siler et al., 2010, 2011, 2012; Brown and Siler, 2014). However, there is growing recognition of the complexity of historical processes that were involved in the diversification of this megadiverse archipelago (see Brown et al., 2013, for a review). For example, there is evidence that older tectonic processes contributed to vertebrate diversification on precursor paleoislands that predates the modern distribution of landmasses in the Philippines (*~*30-5 mya; Jansa et al., 2006; Blackburn et al., 2010; Siler et al., 2012; Brown and Siler, 2014). Additionally, the region’s biodiversity has likely been shaped further by dispersal events from mainland source populations via recognized colonization routes (Diamond and Gilpin, 1983; Brown and Guttman, 2002; Brown and Siler, 2014) and finer-scale isolating mechanisms that led to in situ diversification (Heaney et al., 2011; Linkem et al., 2011; Siler et al., 2011; Hosner et al., 2013). Nonetheless, the question remains: Was climate-driven fragmentation of the islands an important process of speciation?

A “species-pump” model of diversification (Jetz et al., 2004; Fjeldså and Rahbek, 2006; Kozak and Wiens, 2010; Sedano and Burns, 2010; Schoville et al., 2012; Papadopoulou and Knowles, 2015) via repeated vicariance predicts that divergences across taxa that occur on historically connected “island archipelagos” should be clustered around times of historical isolating mechanisms. This model can be relevant to a diversity of structured environments, including deep-ocean (Ricklefs and Bermingham, 2008; Brown et al., 2013; Papadopoulou and Knowles, 2015) and coastal (Papadopoulou and Knowles, 2017; Senczuk et al., 2018) islands, and mountain tops (i.e., “sky” islands; Knowles, 2000, 2001; McCormack et al., 2008) Within the Philippines, this model predicts that divergences among taxa distributed across islands within the same PAIC should be associated with times of rising sea levels that fragmented PAICs into the islands of today. Therefore, if we compare the divergence times of multiple pairs of populations or closely related species that occur on two islands that were connected during glacial periods of lower sea levels, we expect some to be contemporaneous with interglacial fragmentation events. Such patterns of shared divergences would be difficult to explain by other mechanisms, such as over-water dispersal.

Oaks et al. (2013) tested this prediction initially by inferring how many unique divergence times best explained mitochondrial sequence data from 22 pairs of populations from across the Philippines, using a model choice method based on approximate-likelihood Bayesian computation (ABC). However, using simulations, they found that this popular ABC approach was demonstrably sensitive to prior assumptions, with a bias toward over-clustering divergence times, both of which rendered the results difficult to interpret and potentially skewed toward interpretations of simultaneous diversification. Oaks (2014) reanalyzed these data with an ABC method that alleviated these issues, and found that reducing the genetic data to a small number of summary statistics left ABC methods with little information to update prior assumptions.

Here we use comparative genomic data and a new full-likelihood Bayesian method to test the hypothesis that repeated fragmentation of islands was a causal mechanism of vicariant diversification among terrestrial vertebrates in the Philippines. By using all the information in thousands of loci from 16 inter-island pairs of gecko populations, we demonstrate a new method that provides the first robust evaluation of this central tenet of the PAIC model of diversification. Our results support independent diversification among pairs of gecko populations, providing evidence against predictions of the PAIC model of diversification, and underscoring the importance for caution against adhering to overly simplistic models of diversification when studying dynamic and biodiverse regions such as the Philippines.

## Methods

### Sampling

For two genera of geckos, *Cyrtodactylus* and *Gekko*, we sampled individuals from pairs of populations that occur on two different islands. Because the climate-mediated fragmentation of the islands was a relatively recent phenomenon, we selected samples from pairs of localities that were inferred to be closely related, but not necessarily sister, from previous, independent genetic data (Siler et al., 2010, 2012, 2014; Welton et al., 2010a,b). In other words, we avoided pairs that we knew *a priori* diverged well before the connectivity cycles, because these cannot provide insight into whether divergences were clustered *during* these cycles. Furthermore, to avoid complications associated with intra-island population structure, we only used localities where previous genetic data were consistent with the samples being from a single population. We also selected pairs of populations that are independent based on previous phylogenetic estimates (i.e., they do not share any branches in previously estimated phylogenies; Siler et al., 2010, 2012, 2014; Welton et al., 2010a,b).

We also sought to sample pairs that spanned islands connected during glacial periods, as well as islands that were never connected (Figure S1 & S2; Amante and Eakins, 2009; Brown et al., 2013; Spratt and Lisiecki, 2016). We included the latter as “controls.” Because these islands were never connected, the distribution of closely-related populations inhabiting them can only be explained by inter-island dispersal. Thus, it seems reasonable to assume that divergence between these populations was either due to dispersal or an earlier intra-island divergence. Either way, the timing of divergences across islands that were never connected are not expected to be clustered across pairs. These controls are important, given the tendency for previous approaches to this inference problem to over-estimate shared divergences (Oaks et al., 2013, 2014). Finding shared divergence times among pairs for which there is no tenable mechanism for shared divergences will indicate a problem and prevent us from misinterpreting shared divergences among pairs spanning islands that were fragmented as evidence for the PAIC model of vicariant diversification. Applying these criteria, for both genera, we identified eight pairs of populations, including a mix of pairs spanning islands that were never connected, islands that were connected, and islands that were possibly connected during glacial lowstands (Figure 1; Tables 1 & S1).

**Figure 1.**
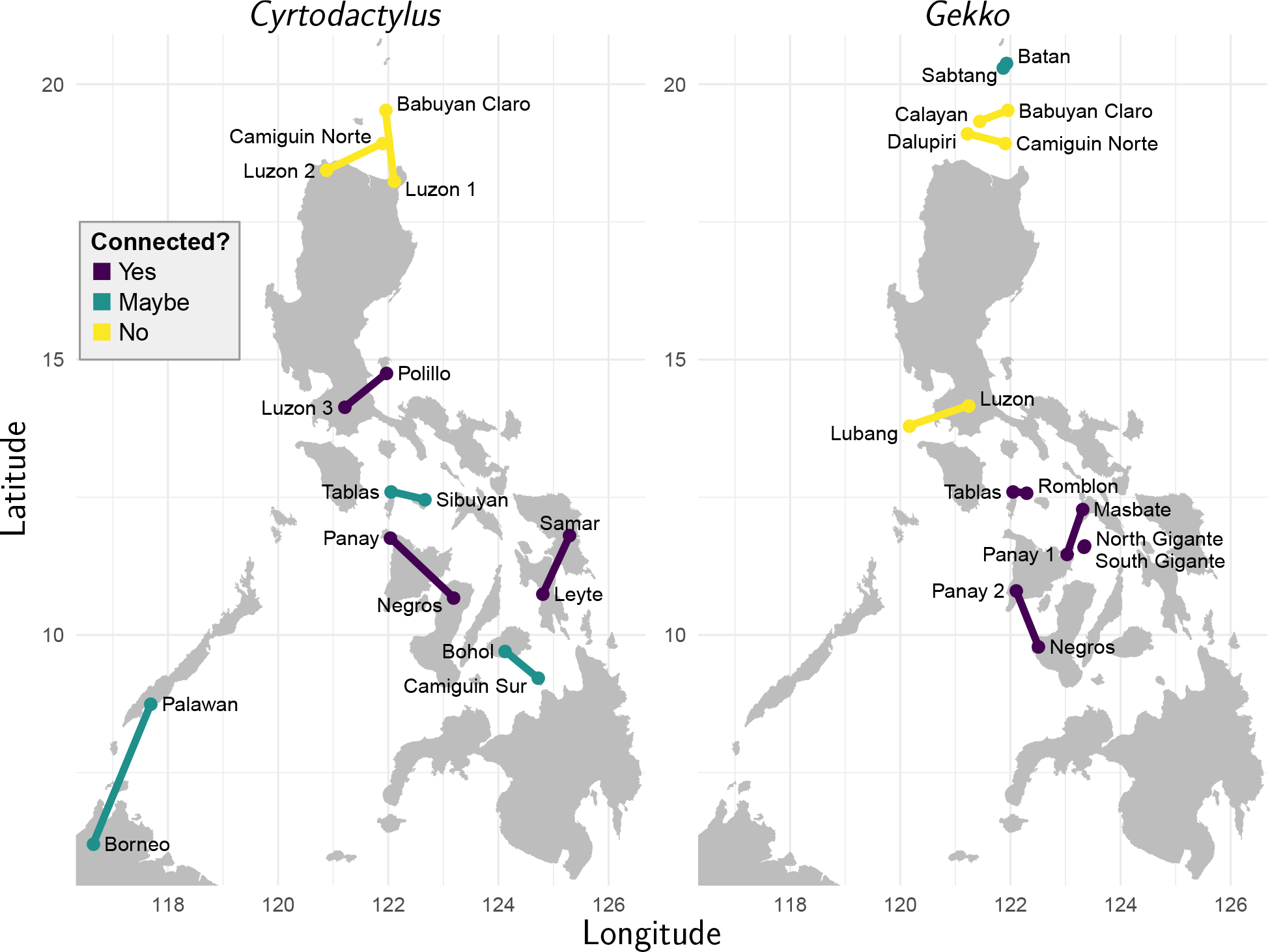
Philippine sampling localities for the eight pairs of *Cyrtodactylus* (left) and *Gekko* (right) populations. Localities for each pair are connected by a line and color-coded (see key) to indicate whether the islands were connected via terrestrial dry land bridges that formed during Pleistocene glacial periods. Figure generated with ggplot2 Version 2.2.1 (Wickham, 2009).

**Table 1.**
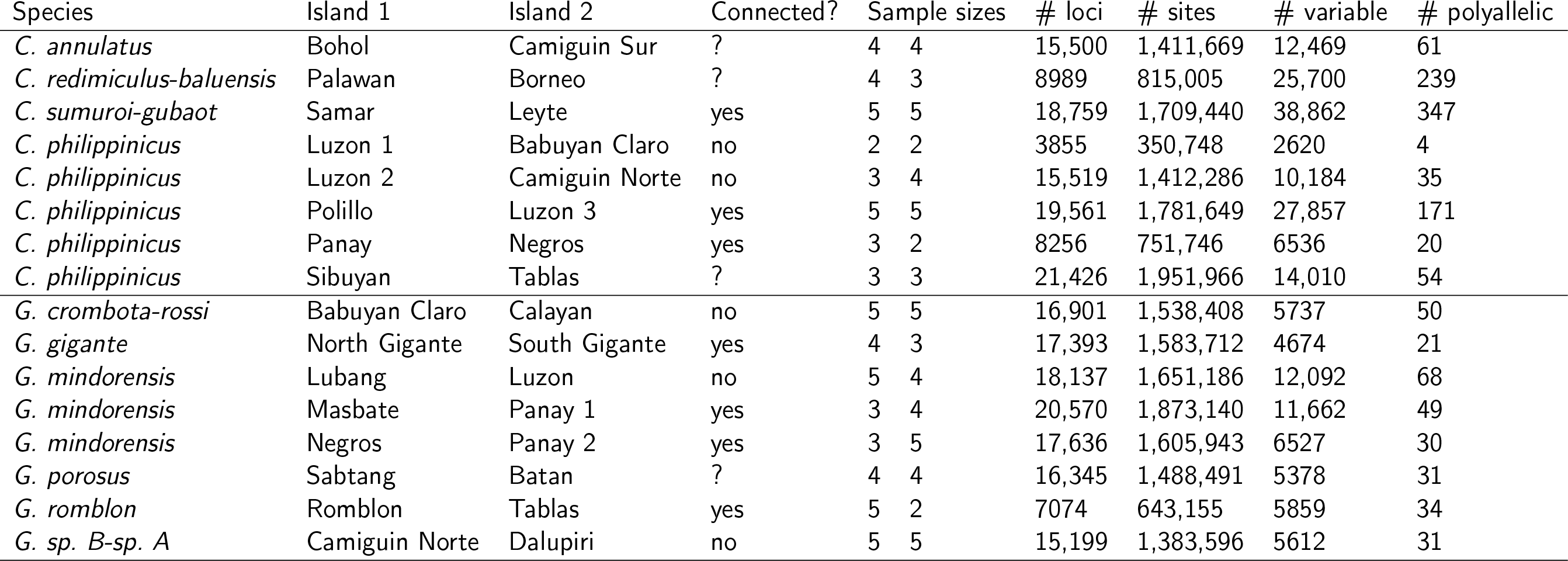
Pairs of *Cyrtodactylus* and *Gekko* populations included in our full-likelihood Bayesian comparative biogeographic analyses (ecoevolity). Each row represents a pair of populations sampled from two islands that either were or were not connected during low sea levels of glacial periods.

The island pairs of Bohol and Camiguin Sur, Palawan and Borneo, Sibuyan and Tablas, and Sabtang and Batan are not believed to have been fully joined during glacial periods. However, even if these pairs of islands did not have a complete land connection, they may have been close enough to permit some limited gene flow given their immediate proximity and the relative dispersal ability of gekkonid lizards. To maximize the number of pairs and thus increase our chances of detecting shared divergences if they occurred, we include these populations in our analyses, but leave their connection status during glacial lowstands as ambiguous (Figure 1; Table 1).

Some of our pairs are currently recognized as different species, whereas others are not (Table 1). Recent taxonomic work on both of these genera of lizards suggests they comprise many more species than previously recognized, with more revisions necessary (Brown et al., 2009; Linkem et al., 2010b; Siler et al., 2010; Welton et al., 2010a,b; Brown et al., 2011; Grismer et al., 2018b,a). Over-water dispersal events are necessary to explain the existence of these populations on oceanic islands. However, these events are likely too random and rare to contribute meaningful levels of gene flow between islands. Therefore, given the insularity of these populations, we assume that none of them are experiencing migration during interglacial periods, like today. Thus, all of the populations we sampled are likely independent evolutionary lineages, regardless of whether they are currently recognized by taxonomists as distinct species.

### Genomic library preparation and sequencing

We extracted DNA from tissue using the guanidine thiocyanate method described by Esselstyn et al. (2008). We diluted the extracted DNA for each individual to a concentration of 5 ng/*μ*L based on the initial concentration measured with a Qubit 2.0 Fluorometer. We generated three restriction-site associated DNA sequence (RADseq) libraries, each with 96 individuals, using the multiplexed shotgun genotyping (MSG) protocol of Andolfatto et. al. (Andolfatto et al., 2011). Following digestion of 50 ng of DNA with the NdeI restriction enzyme, we ligated each sample to one of 96 adaptors with a unique six base-pair barcode. After pooling the 96 samples together, we selected 250-300bp fragments to remain in the library using a Pippen Prep. For each pool of 96 size-selected samples, we performed eight separate polymerase chain reactions for 14 cycles (PCR) using Phusion High-Fidelity PCR Master Mix (NEB Biolabs) and primers that bind to common regions in the adaptors. Following PCR, we did two rounds of AMPure XP bead cleanup (Beckman Coulter, Inc.) using a 0.8 bead volume to sample ratio. Each library was sequenced in one lane of an Illumina Hiseq 2500 high-output run, with single-end 100bp reads. We provide information on all of the individuals included in the three RADseq libraries in Table S1, a subset of which were included in the population pairs we analyzed for this study (Table 1 & S2). We deposited the demultiplexed, raw sequence reads into the NCBI Sequence Read Archive (Bioproject PRJNA486413, SRA Study SRP158258), and the assembled data matrices are available in our project repository (https://github.com/phyletica/gekgo).

### Data assembly

We used ipyrad version 0.7.13 (Eaton, 2017) to demultiplex and assemble the raw RADseq reads into loci. To maximize the number of assembled loci, we de novo assembled the reads separately for each pair of populations. All of the scripts and ipyrad parameters files we used to assemble the data are available in our project repository (https://github.com/phyletica/gekgo), and the ipyrad settings are listed in Table S3.

### Testing for shared divergences

We approach the inference of temporally clustered divergences as a problem of model choice. Our goal is to treat the number of divergence events shared (or not) among the pairs of populations, and the assignment of the pairs to those events, as random variables to be estimated from the aligned sequence data. For eight pairs, there are 4,140 possible divergence models (i.e., there are 4,140 ways to partition the eight pairs to *k* = 1, 2, …, 8 divergence events; Bell, 1934; Oaks, 2014, 2018). Although divergences caused by sea-level rise would not happen simultaneously, we expect that on a timescale of the lizards’ mutation rate, treating them as simultaneous should be a better explanation of data generated by such a process than treating them as independent.

Given the large number of models, and our goal of making probability statements about them, we used a Bayesian model-averaging approach. Specifically, we used the full-likelihood Bayesian comparative biogeography method implemented in the software package `ecoevolity` version 0.1.0 (commit b9f34c8) (Oaks, 2018). This method models each pair of populations as a two-tipped “species” tree, with an unknown, constant population size along each of the three branches, and an unknown time of divergence, after which there is no migration. This method can directly estimate the likelihood of values of these unknown parameters from orthologous biallelic characters by analytically integrating over all possible gene trees and mutational histories (Bryant et al., 2012; Oaks, 2018). Within this full-likelihood framework, this method uses a Dirichlet process prior on the assignment of our pairs to an unknown number of divergence times. The Dirichlet process is specified by a (1) concentration parameter, *α*, which determines how probable it is for pairs to share the same divergence event, *a priori*, and (2) base distribution, which serves as the prior on the unique divergence times.

Importantly, because the pairs of populations are modeled as disconnected species trees, the relative rates of mutation among the pairs is not identifiable. This requires us to make informative prior assumptions about the relative rates of mutation among the pairs. Because *Cyrtodactylus* and *Gekko* are deeply divergent (> 80 mya; Gamble et al., 2011), and nothing is known about their relative rates of mutation, we analyzed the two genera separately. Within each genus, the populations are all closely related (Siler et al., 2010, 2012, 2014; Welton et al., 2010a,b) allowing us to make the simplifying assumption that all pairs *within* each genus share the same rate of mutation. Furthermore, we set the rate to one so that effective population sizes and time are scaled by the mutation rate, and thus time is in expected substitutions per site.

Based on previous data (Siler et al., 2010; Welton et al., 2010a,b) we assumed a prior on divergence times of *τ ~* Exponential(mean = 0.005) for our eight pairs of *Cyrtodactylus* populations, in units of substitutions per site. To explore the sensitivity of our results to this assumption, we also tried a prior on divergence times of *τ ~* Exponential(mean = 0.05). Based on previous data (Siler et al., 2012, 2014), we assumed a divergence-time prior of *τ ~* Exponential(mean = 0.0005) for our eight pairs of *Gekko* populations, in units of substitutions per site. To explore the sensitivity of our results to this assumption, we also tried priors of Exponential(mean = 0.005) and Exponential(mean = 0.05) on the *Gekko* divergence times.

For the concentration parameter of the Dirichlet process, we assumed a hyperprior of *α ~* Gamma(1.1, 56.1) for both genera. This places approximately half of the prior probability on the model with no shared divergences (*k* = 8). By placing most of the prior probability on the model of independent divergences, if we find posterior support for shared divergences, we can be more confident it is being driven by the data, as opposed to the prior on divergence times penalizing additional divergence-time parameters (Jeffreys, 1939; Lindley, 1957; Oaks et al., 2013, 2014). To explore the sensitivity of our results to this assumption, we also tried a hyperprior of *α ~* Gamma(1.5, 3.13) and *α ~* Gamma(0.5, 1.31). The former corresponds to a prior mean number of divergence events of five, whereas the latter places 50% of the prior probability on the single divergence (*k* = 1) model.

For all analyses of both the *Cyrtodactylus* and *Gekko* data, we assumed equal mutation rates among the pairs, a prior distribution of Gamma(shape = 4.0, mean = 0.004) on the effective size of the populations scaled by the mutation rate (*N*_*e*_*μ*), and a prior distribution of Gamma(shape = 100, mean = 1) on the relative effective size of the ancestral population (relative to the mean size of the two descendant populations).

The model implemented in `ecoevolity` assumes each character is unlinked (i.e., evolved along a gene tree that is independent conditional on the population tree). Data that satisfy this assumption include single-nucleotide polymorphisms (SNPs) that are well-spaced across the genome. However, by analyzing simulated data, Oaks (2018) showed the method performs better when all linked sites are used than when data are excluded to avoid violating the assumption of unlinked sites. We simulate data sets based on our gekkonid data to confirm these results hold for our sampling design (see below). Furthermore, when analyzing the RADseq from three of the *Gekko* population pairs we are analyzing here, Oaks (2018) found biologically unrealistic estimates of divergence times and population sizes when only unlinked variable sites (i.e., SNPs) were analyzed. Using additional simulations, Oaks (2018) found these unrealistic estimates were likely due to data-acquisition biases, which are known to be common in alignments from reduced-representation genomic libraries (Harvey et al., 2015; Linck and Battey, 2019). Oaks (2018) found that using all of the sites, rather than only SNPs, greatly improved the robustness of these parameter estimates to such acquisition biases. Considering all these findings, we are confident in the inclusion of all sites of our RADseq loci in the `ecoevolity` analyses. Given that all sites were used, the likelihood computed in `ecoevolity` was not conditioned on only sampling variable characters (Oaks, 2018).

The model implemented in `ecoevolity` is also restricted to characters with two possible states (biallelic). Thus, for sites with three or more nucleotides (hereafter referred to as polyallelic sites), we compared how sensitive our results were to two different strategies: (1) Removing polyallelic sites, and (2) recoding the sites as biallelic by coding each state as either having the first nucleotide in the alignment or a different nucleotide. We assumed the biallelic equivalent of a Jukes-Cantor model of character substitution (Jukes and Cantor, 1969) so that our results are not sensitive to how nucleotides are coded as binary (Oaks, 2018).

For each analysis, we ran 10 independent Markov chain Monte Carlo (MCMC; Metropolis et al., 1953; Hastings, 1970) chains for 150,000 generations, sampling every 100th generation. We assessed convergence and mixing of the chains by inspecting the potential scale reduction factor (PSRF; the square root of Equation 1.1 in Brooks and Gelman, 1998) and effective sample size (Gong and Flegal, 2016) of the log likelihood and all continuous parameters using the `pyco-sumchains` tool of `pycoevolity`. We also visually inspected the trace of the log likelihood and parameters over generations with the program Tracer version 1.6 (Rambaut et al., 2014).

### Vetting our sampling and methodology

#### Simulation-based assessment of ecoevolity conditional on our sampling

Oaks (2018) tested the method implemented in `ecoevolity` using simulated data. However, our gekkonid RADseq data differ from the simulation conditions used by Oaks (2018) in a number of ways. For example, we have fewer individuals sampled from most of our populations than the five simulated by Oaks (2018), and the number of loci and sites vary dramatically among our pairs of populations (Table 1).

To assess how well `ecoevolity` is able to infer shared divergences based on our sampling design, we implemented new simulation options in the `simcoevolity tool` within the `ecoevolity` software package. Our modifications allow us to simulate data sets that exactly match the sampling scheme of our gekkonid data. Specifically, we simulate data sets that match our empirical data in terms of

1. the number of loci for each pair of populations,
2. the number of sites within each locus, and
3. the number of gene copies sampled for each site (i.e., the same patterns of missing data).

We assume all loci are effectively unlinked with no intralocus recombination (i.e., all the sites of a locus evolved along the same gene tree that is independent of the other loci, conditional on the population history). Our simulator allows us to sample all sites from each locus, or only a maximum of one variable site per locus. The former violates the assumption of the model implemented in `ecoevolity` that all sites are effectively unlinked, whereas the latter avoids this model violation at the cost of excluding data.

For each genus, we simulated 500 fully sampled data sets and 500 data sets with, at most, one SNP sampled per locus. For each simulation, the divergence model and all parameter values were drawn from the same distributions we used as priors in our empirical analyses described above. Specifically, the concentration parameter of the Dirichlet process was drawn from a gamma distribution of Gamma(1.5, 3.13), and the time of each divergence event was distributed as Exponential(mean = 0.005) and Exponential(mean = 0.0005) for *Cyrtodactylus* and *Gekko*, respectively. When analyzing each simulated dataset with `ecoevolity`, we used these same distributions as priors and ran four independent MCMC chains for 150,000 generations, sampling every 100th generation. After ignoring the first 501 “burn-in” samples from each chain (including the initial state), we collected 4,000 MCMC samples for each analysis.

#### Data partitioning to evaluate performance of ecoevolity

With our simulations, we can assess how well `ecoevolity` infers shared divergences we know to be true. However, the simulated data sets are undoubtedly simpler than our empirical data, without any model violations (barring the linked sites within loci) introducing variation. Thus, we took another approach using our gekkonid RADseq data directly. We split the 21,426 loci randomly from our population pair with the largest number of sampled loci (*C. philippinicus* from the islands of Sibuyan and Tablas) into two subsets of 10,713 loci. We then reanalyzed the data with `ecoevolity` using the methods described above, but treating the two subsets of loci as separate population pairs. If `ecoevolity` can reliably detect a shared divergence event, it should infer that the two sets of loci from the same pair of populations did indeed co-diverge.

## Results

### Data collection and MCMC convergence

Table 1 summarizes the number of individuals sampled for each pair of islands, along with the number of assembled loci, and the number of total, variable, and polyallelic characters. The nucleotide diversity within and between each pair of populations is provided in Table S4. The 10 independent MCMC chains of all our `ecoevolity` analyses appeared to have converged almost immediately. We conservatively removed the first 101 samples, leaving 1,400 samples from each chain (14,000 samples for each analysis). With the first 101 samples removed, across all our analyses, all ESS values were greater than 2,000, and all PSRF values were less than 1.005.

### Testing for shared divergences

#### *Cyrtodactylus* population pairs

For *Cyrtodactylus*, our `ecoevolity` results support the model of no shared divergences, i.e., all eight pairs of populations diverged independently (Figures 2 & 3). This support is consistent across all three priors on the concentration parameter of the Dirichlet process (Figures S3 & S4). The support is also consistent across both priors on divergence times and whether polyallelic sites are recoded or removed (Figures S5 & S6). Estimates of effective population sizes are also very robust to priors on *α* and *τ*, and whether polyallelic sites are recoded or removed (Figures S7 & S8).

**Figure 2.**
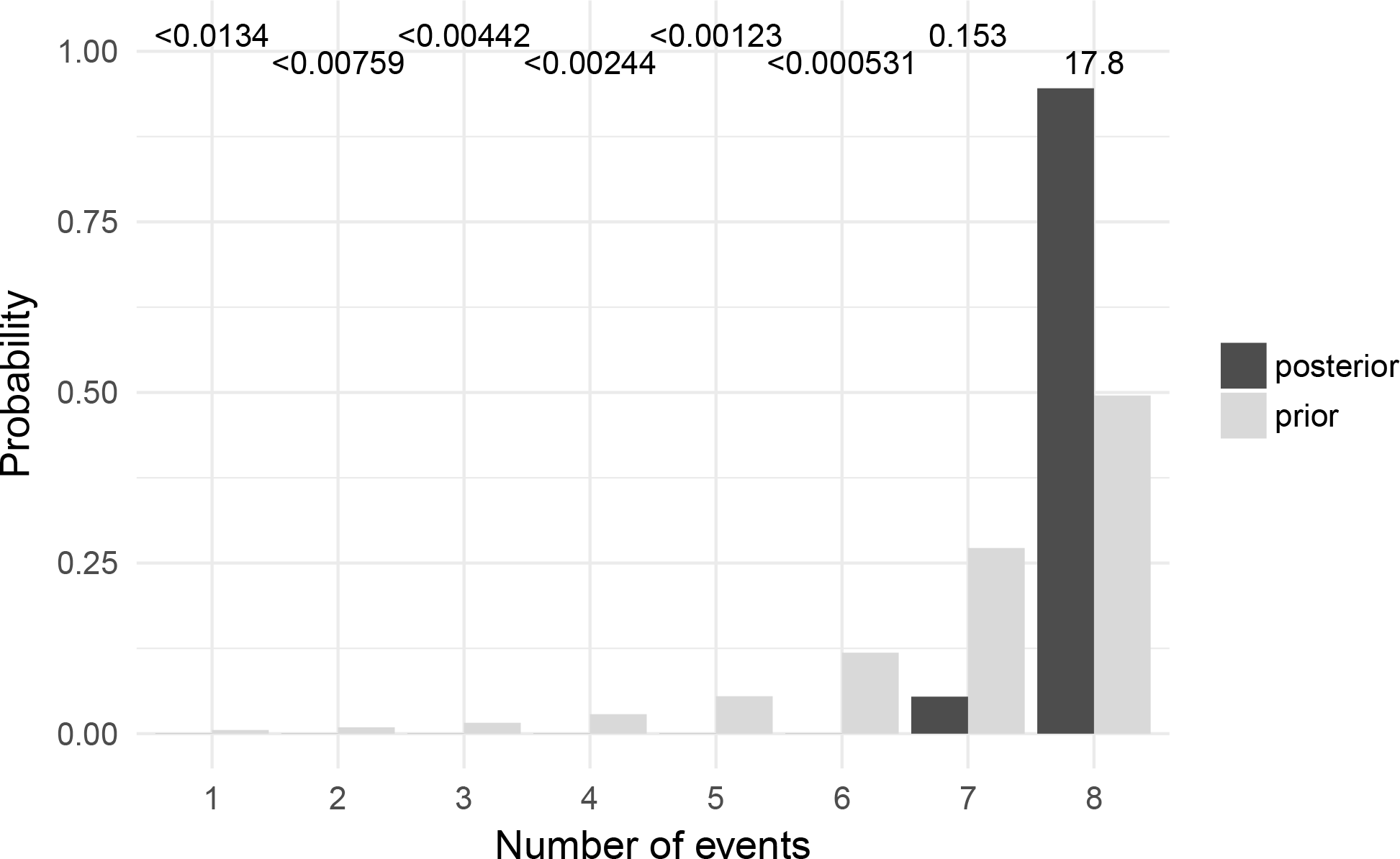
Approximate prior (light bars) and posterior (dark bars) probabilities of numbers of divergence events across pairs of *Cyrtodactylus* populations. Bayes factors for each number of divergence times is given above the corresponding bars. Each Bayes factor compares the corresponding number of events to all other possible numbers of divergence events. Figure generated with ggplot2 Version 2.2.1 (Wickham, 2009).

**Figure 3.**
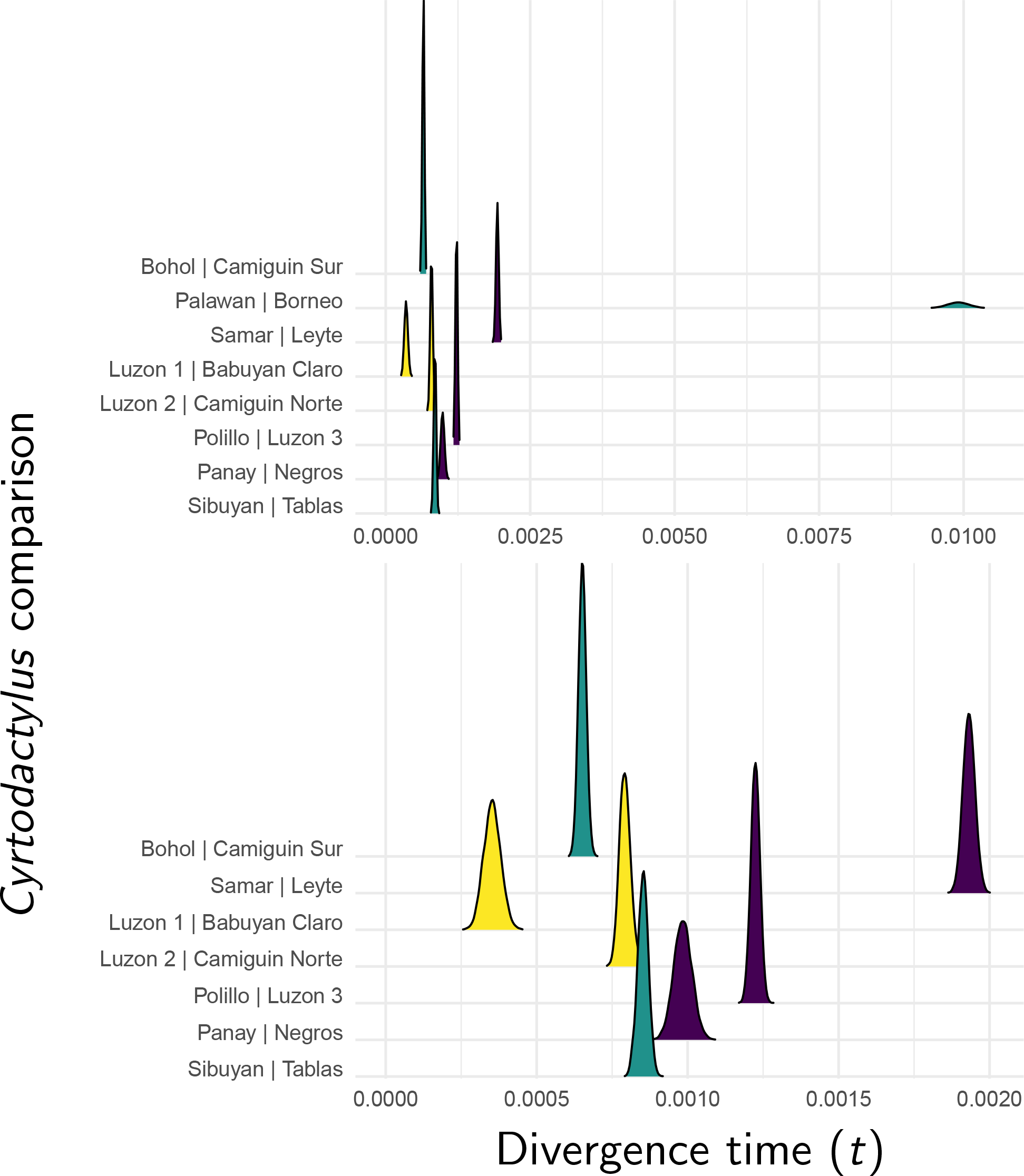
Approximate marginal posterior densities of divergence times (in expected substitutions per site) for each pair of *Cyrtodactylus* populations. The density plot of each pair is color-coded to indicate whether the islands were connected during glacial periods (Fig. 1). The top plot shows all eight pairs of populations, whereas the bottom plot excludes the pair of *C. redimiculus* and *C. baluensis* from Palawan and Borneo. Figure generated with ggridges Version 0.4.1 (Wilke, 2018) and ggplot2 Version 2.2.1 (Wickham, 2009).

#### *Gekko* population pairs

For *Gekko*, posterior probabilities weakly support no shared divergences, but Bayes factors weakly support seven divergence events across the eight pairs (Figure 4), suggesting a possible shared divergence between *G. mindorensis* on the islands of Panay and Masbate and *G. porosus* on the islands of Sabtang and Batan (Figure 5). Under the intermediate prior on the concentration parameter, support increases for this shared divergence (Figures S9 & S10). Under the prior that puts most of the probability on one shared event, posterior probabilities prefer six divergences (Figure S9) with another shared divergence between *G. crombota* and *G. rossi* on the islands of Babuyan Claro and Calayan and *G. romblon* on the islands of Romblon and Tablas (Figure S10); however, Bayes factors still prefer seven divergences. Similarly, as the prior on divergence times becomes more diffuse, the results shift from ambiguity between seven or eight divergence events, to ambiguity between six or seven events, to strong support for six events, with the same island pairs sharing divergences (Figures S11 & S12).

**Figure 4.**
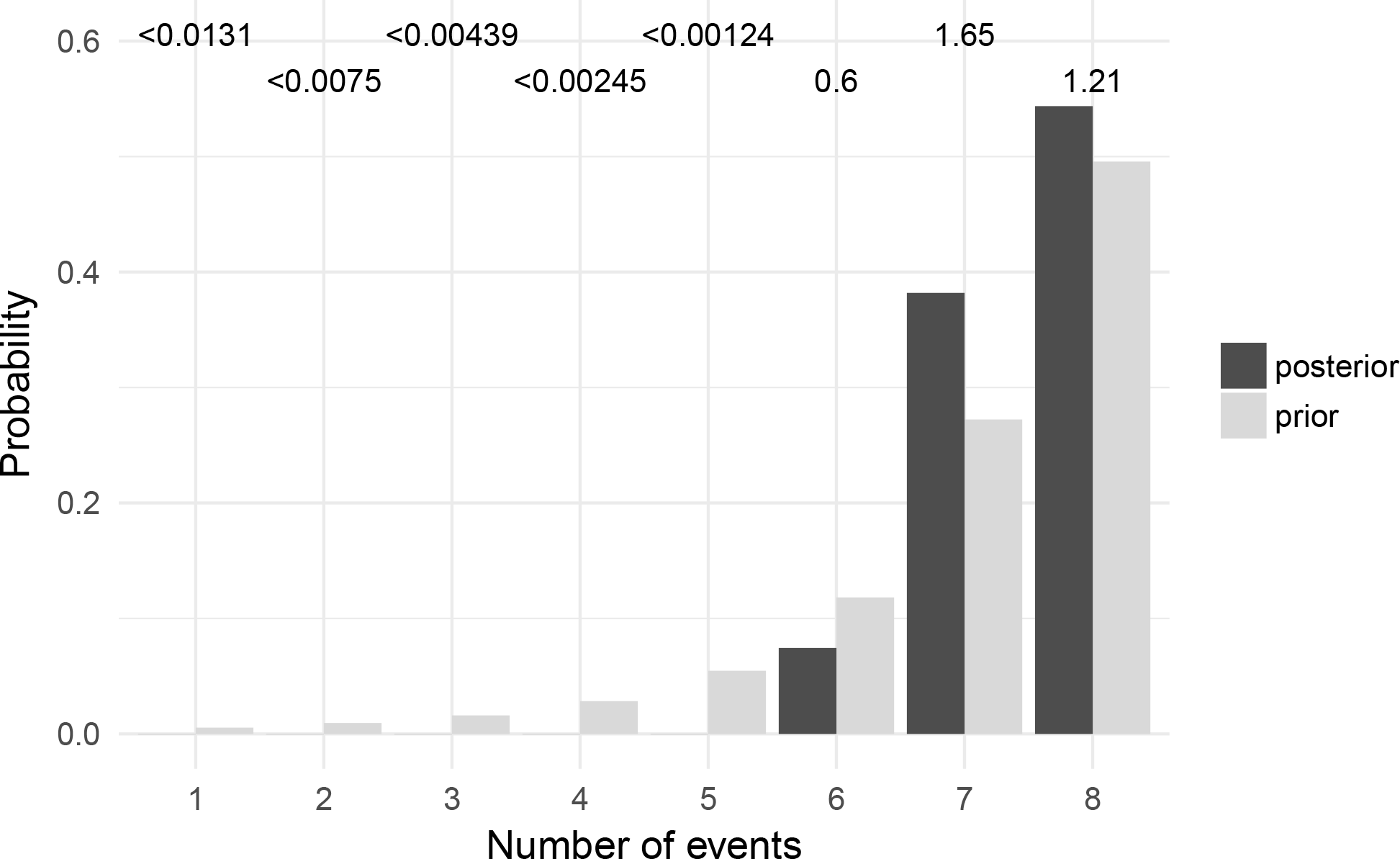
Approximate prior (light bars) and posterior (dark bars) probabilities of numbers of divergence events across pairs of *Gekko* populations. Bayes factors for each number of divergence times is given above the corresponding bars. Each Bayes factor compares the corresponding number of events to all other possible numbers of divergence events. Figure generated with ggplot2 Version 2.2.1 (Wickham, 2009).

**Figure 5.**
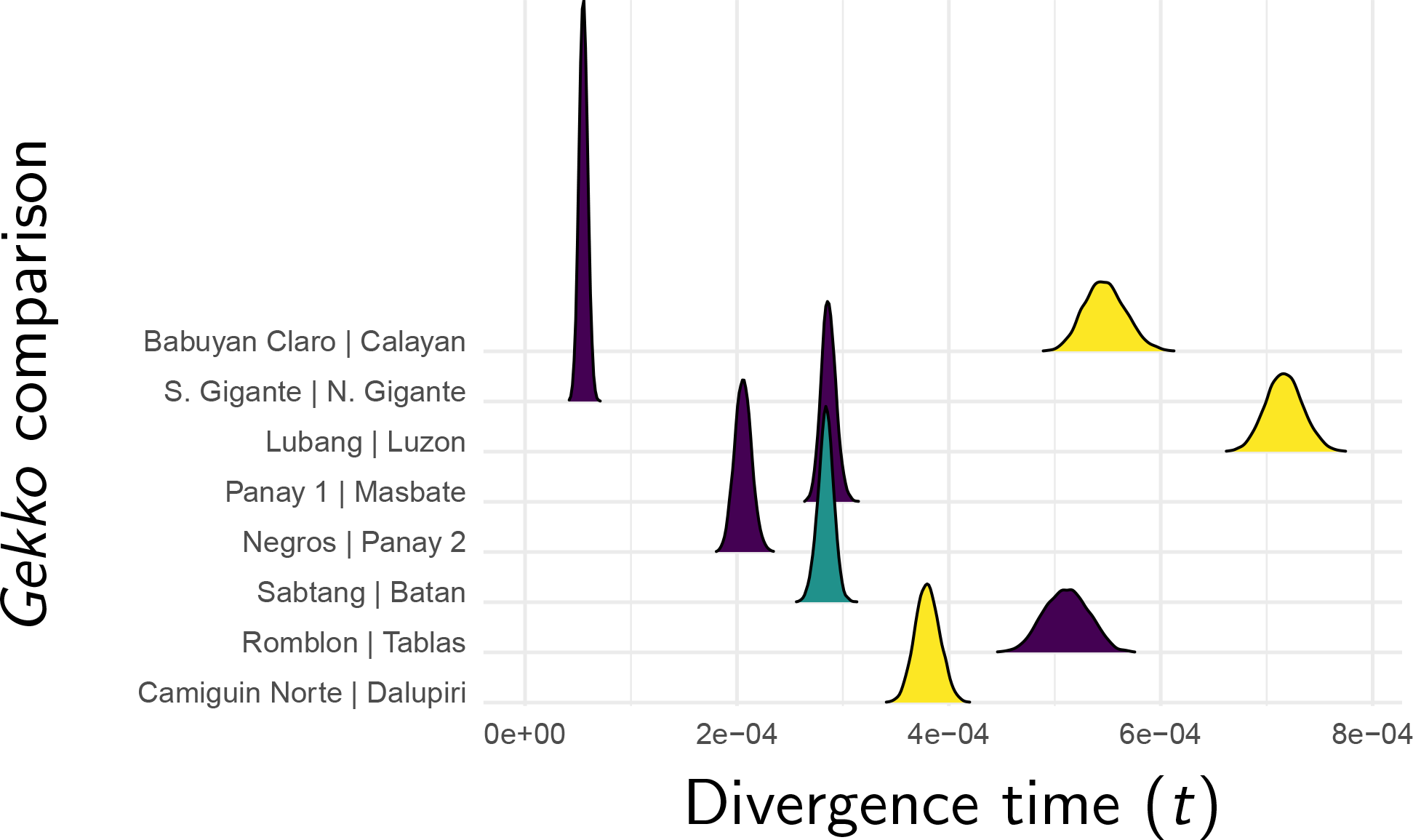
Approximate marginal posterior densities of divergence times (in expected substitutions per site) for each pair of *Gekko* populations. The density plot of each pair is color-coded to indicate whether the islands were connected during glacial periods (Fig. 1). Figure generated with ggridges Version 0.4.1 (Wilke, 2018) and ggplot2 Version 2.2.1 (Wickham, 2009).

As with *Cyrtodactylus*, the estimates of divergence times are robust to whether polyallelic sites are recoded or removed (Figure S12), and population size estimates are robust to priors on *α* and *τ*, and whether polyallelic sites are recoded or removed (Figures S13 & S14).

### Simulation results

The results from analyses of the simulated data sets show our gekkonid RADseq data are sufficient for `ecoevolity` to accurately estimate the timing (Figure 6) and number (Figure 7) of divergence events, and the effective sizes of the ancestral (Figure S15) and descendant populations (Figure S16). Consistent with Oaks (2018), we also find that estimation accuracy and precision are much better when all sites are analyzed rather than only unlinked SNPs.

**Figure 6.**
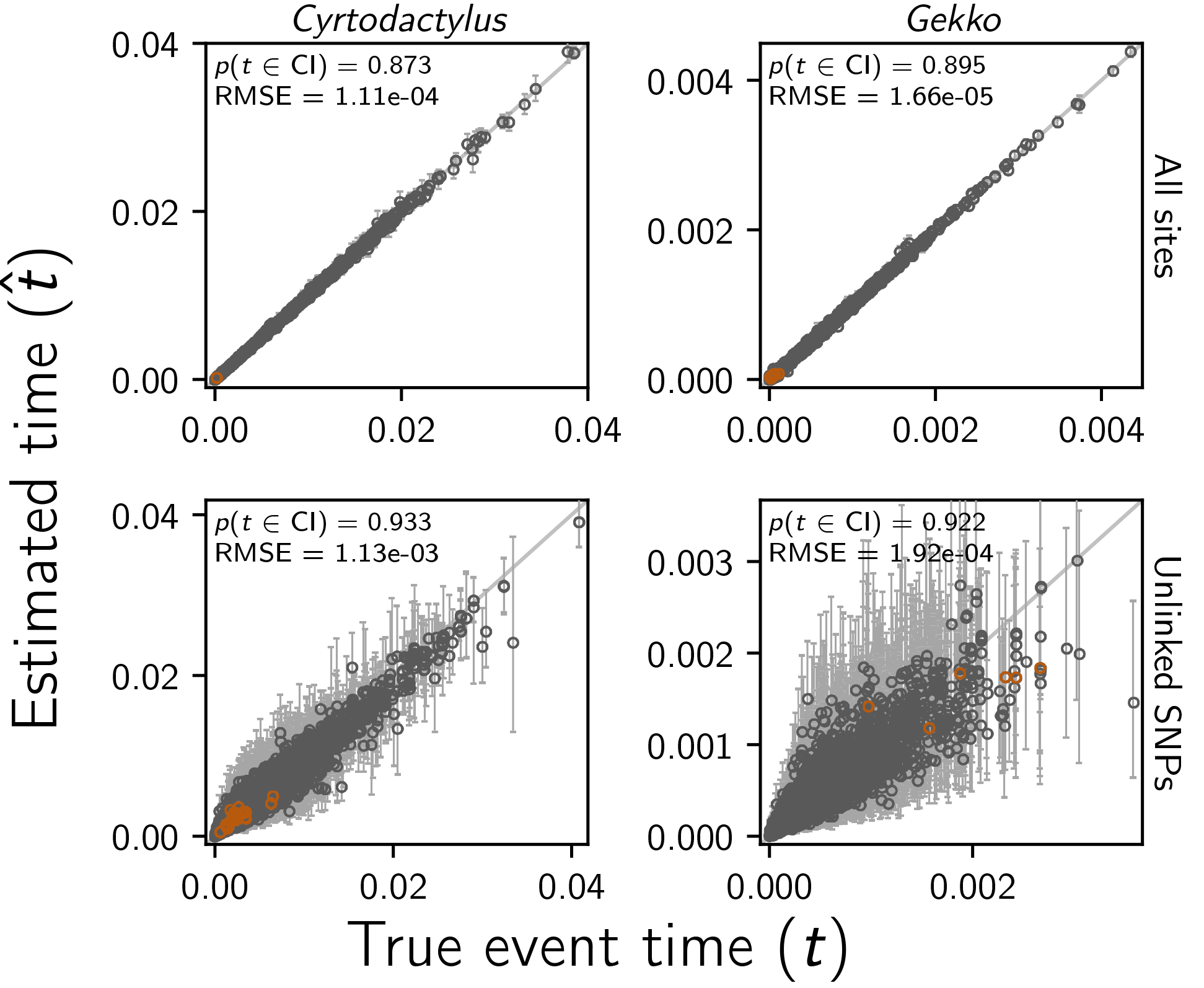
The accuracy and precision of `ecoevolity` divergence-time estimates (in units of expected subsitutions per site) when applied to data simulated to match our *Cyrtodactylus* (left) and *Gekko* (right) RADseq data sets with all sites (top) or only one SNP per locus (bottom). Each circle and associated error bars represents the posterior mean and 95% credible interval for the time that a pair of populations diverged. Estimates for which the potential-scale reduction factor was greater than 1.2 (Brooks and Gelman, 1998) are highlighted in orange. Each plot consists of 4,000 estimates—500 simulated data sets, each with eight pairs of populations. For each plot, the root-mean-square error (RMSE) and the proportion of estimates for which the 95% credible interval contained the true value—*p*(*t* ∈ CI)—is given. Figure generated with matplotlib Version 2.0.0 (Hunter, 2007).

**Figure 7.**
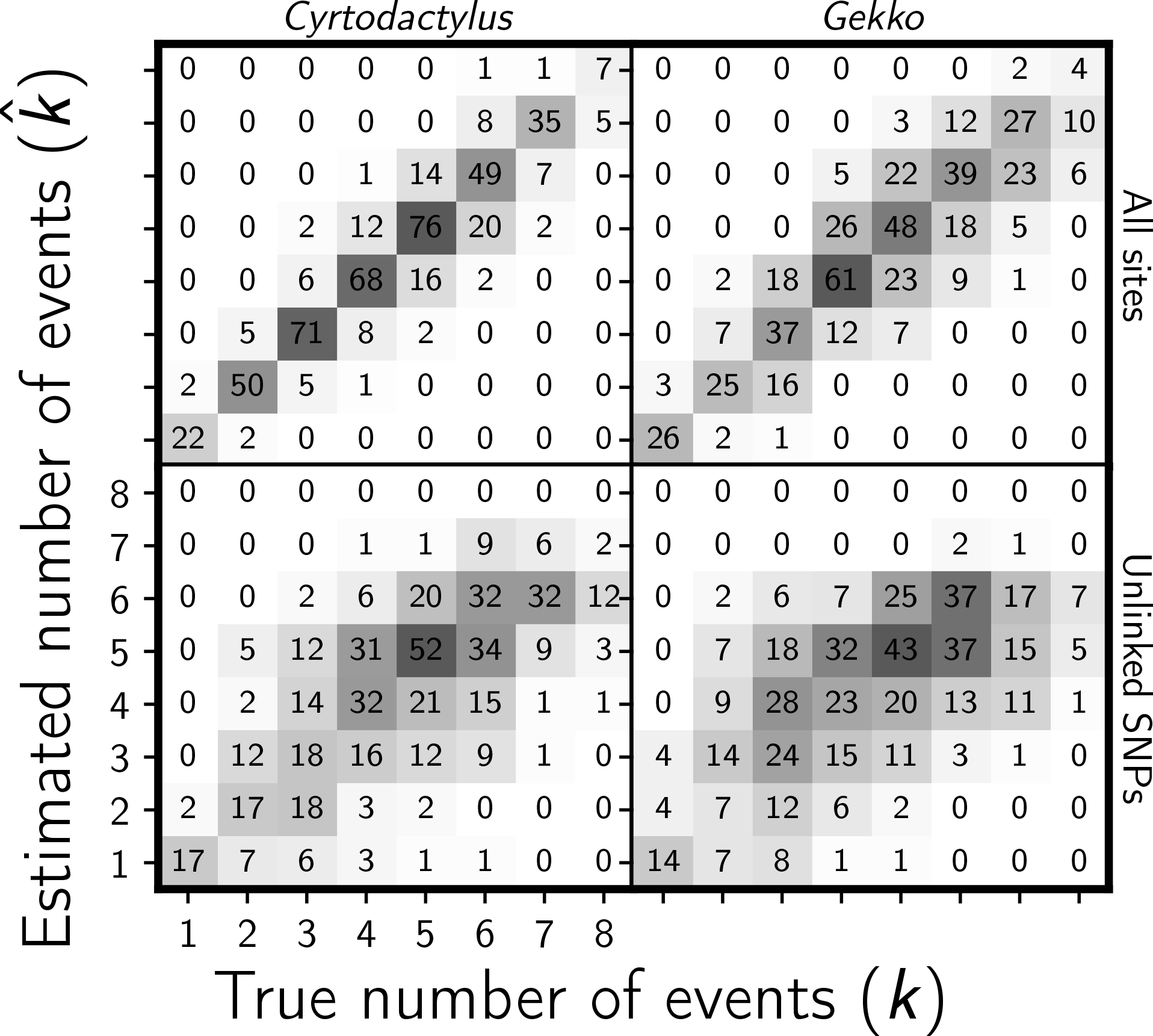
The accuracy of `ecoevolity` to estimate the number of divergence events when applied to data simulated to match our *Cyrtodactylus* (left) and *Gekko* (right) RADseq data sets with all sites (top) or only one SNP per locus (bottom). Each plot illustrates the results of the analyses of 500 simulated data sets, each with eight pairs of populations. The number of simulation replicates that fall within each possible cell of true versus estimated numbers of events is shown, and cells with more replicates are shaded darker. Figure generated with matplotlib Version 2.0.0 (Hunter, 2007).

### Data partitioning to vet ecoevolity

When we reanalyzed our *Cyrtodactylus* data with the loci from the pair of *C. philippinicus* populations from the islands of Sibuyan and Tablas randomly split into two sets and treated as separate comparisons, the Bayes factors approximated by `ecoevolity` strongly support (Jefferys, 1961) that the two subsets of loci codiverged. The posterior odds of them codiverging was 72.83 (posterior probability 0.963) and 144.81 (posterior prorbability 0.889) times greater than the prior odds when the hyperprior on the concentration parameter of the Dirichlet process was Gamma(1.5, 3.13) and Gamma(1.1, 56.1), respectively. We performed these calcuations with the sumcoevolity tool of the `ecoevolity` software package, using one million simulations under the Dirichlet process to approximate the prior odds.

## Discussion

### Is there evidence for shared divergences among *Gekko* pairs?

Under some of the priors we explored, there is support for two possible shared divergences among the pairs of *Gekko* populations: (1) *G. mindorensis* on the islands of Panay and Masbate and *G. porosus* on the islands of Sabtang and Batan, and (2) *G. crombota* and *G. rossi* on the islands of Babuyan Claro and Calayan and *G. romblon* on the islands of Romblon and Tablas. The islands of Babuyan Claro and Calayan were never connected, and we only inferred support for the second shared divergence under the most extreme priors on *α* and *τ* that are expected to favor shared divergences (Figures S9, S10, S11, & S12). Thus, support for the second shared divergence scenario is likely an artifact of prior sensitivity. However, the weak support for a shared divergence between *G. mindorensis* on the islands of Panay and Masbate and *G. porosus* on the islands of Sabtang and Batan under more reasonable priors is interesting because both pairs of islands were either connected or potentially close enough during glacial periods to allow gene flow.

Under the priors we initially chose as appropriate (as opposed to those used to assess prior sensitivity), the posterior probability that the Panay-Masbate and Sabtang-Batan pairs co-diverged is 0.385. To evaluate support for this co-divergence, we could calculate a Bayes factor using the prior probability that any two pairs share the same divergence time, which is approximately 1.66 in favor of the co-divergence (Figure 4). However, this would not be appropriate, because we did not identify the Panay-Masbate and Sabtang-Batan pairs of interest *a priori*, but rather our attention was drawn to these pairs based on the posterior results. Thus, the probability that *any* two pairs share the same divergence is no longer the appropriate prior probability for our Bayes factor calculation. Rather, we need to consider the prior probability that the two pairs with the most similar divergence times share the same divergence. To get this prior probability, we can take advantage of the fact that this condition is met anytime the number of divergence events is less than eight. Thus, the prior probability that the two pairs with the most similar divergence times share the same divergence is equal to one minus the prior probability that all eight pairs diverge independently. Under our prior on the concentration parameter of Gamma(1.1, 56.1), this prior probability is approximately 0.5. Therefore, our posterior probability for the co-divergence between the Panay-Masbate and Sabtang-Batan pairs is actually *less* than the prior probability, resulting in a weak Bayes factor of approximately 1.6 in support *against* the co-divergence. Based on probability theory, we should favor the explanation that all eight pairs of *Gekko* populations diverged independently.

### Caveats and implications for the PAIC diversification model

#### Limited numbers of comparisons

For each genus we sampled 5 or 6 pairs of populations that span two islands that were connected (or nearly so) by terrestrial dry land bridges during Pleistocene glacial periods (Brown and Diesmos, 2009; Brown et al., 2013). The connectivity between each pair of islands was likely fragmented by rising sea levels six or more times over the past three million years (Figures S1 & S2; Rohling et al., 1998; Siddall et al., 2003; Spratt and Lisiecki, 2016; Amante and Eakins, 2009). Given that, for each genus, we have fewer pairs than the number of times the islands were fragmented, the support we found for independent divergence times among the pairs we analyzed does not obviate all correlates of the PAIC diversification model; our pairs could have diverged at different fragmentation events. Comparative genomic data from more pairs of populations would be necessary to explore this possibility. Nonetheless, with seven pairs (11 including ambiguous island connections), we find no support for shared divergences, suggesting that, at the very least, climate-mediated vicariance is not the primary mode of population divergence in these insular gekkonids.

#### Variation in fragmentation times among island pairs

Another possibility is that some of our pairs of populations diverged during the same interglacial period, but the time when gene flow was cut off by rising sea levels was different enough to be estimated as separate divergences in `ecoevolity`. Based on bathymetry data (Amante and Eakins, 2009), all of the previously connected pairs of islands we sampled (Figure 1) were connected when sea levels were 5-15m below current levels (Figure S1). Based on these bathymetry data and the sea level projections of Spratt and Lisiecki (2016), the timing of fragmentation among these pairs of islands would have differed by less than 3,000 years during the last two interglacial periods (Figure S2). If we assume a rate of mutation an order of magnitude faster than that estimated by Siler et al. (2012) for the phosducin gene of Philippine *Gekko* (1.18×10^−9^ substitutions per site per year), we would not expect to see a difference in divergence times greater than 3.5*×*10^−6^ substitutions per site between two pairs that diverged during the same interglacial. This is likely an over-estimate, given that both the number of years between island separations and the mutation rate are toward the upper end of plausible.

It seems reasonable to assume that the difference in divergence times between *G. mindorensis* on the islands of Panay and Masbate and *G. porosus* on the islands of Sabtang and Batan is close to the minimum resolution of `ecoevolity` given our data; there is little posterior variance in the divergence times for these pairs (Figure 5), and we note posterior uncertainty in whether these pairs co-diverged or not (Figure 4). The posterior mean absolute difference in divergence time between these pairs, conditional on them not co-diverging, is 9.66*×*10_−6_ substitutions per site. This is more than 2.7 times larger than our maximum expected divergence within an interglacial cycle of 3.5×10^−6^ substitutions per site, suggesting that `ecoevolity` would not have the temporal resolution given our data to distinguish the divergence times of two pairs that diverged during the same interglacial fragmentation event. Also, among all remaining pairs, we found support for distinct divergences with much larger differences than expected within an interglacial period. Thus, it seems unlikely that variation in island separation times within interglacial periods explains the variation in divergence times we see across the pairs of gekkonid populations.

Nonetheless, it would be ideal to sample pairs of populations that are co-distributed across the same pair of islands so that we know the fragmentation occurred at the same time. However, doing so comes with the inherent trade-off of having to compare more distantly related taxa. For example, we could sample pairs of *Cyrtodactylus* and *Gekko* populations that span the same islands, but not multiple pairs within each genus. This is important, because to compare divergence times among taxa, we need to make strong assumptions about their relative rates of mutation. Without information about mutation rates, we can assume equal rates across the comparisons, as we did here, but this assumption becomes much more questionable as the taxa we wish to compare are more distantly related from one another. Given the variation among island fragmentation times is small (< 3,000 years) relative to evolutionary timescales, we feel this trade-off is more desirable than making simplifying assumptions about relative mutation rates among distantly related taxa. However, if information about relative rates of mutation among taxa that span the same islands can be brought to bare, an analysis of such a system would provide a strong and complementry empirical test of the PAIC model.

#### Assumptions about mutation rates

As discussed above, we made the simplifying assumption that mutation rates were equal across pairs and constant through time. To minimize the impact of violations of this assumption, we analyzed the gekkonid genera separately, each of which comprise species that are closely related relative to other comparative phylogeographic studies that have made this assumption (Hickerson et al., 2006; Leaché et al., 2007; Plouviez et al., 2009; Voje et al., 2009; Barber and Klicka, 2010; Daza et al., 2010; Chan et al., 2011; Huang et al., 2011; Oaks et al., 2013; Stone et al., 2012; Smith et al., 2014). For example, all of the populations of *Gekko* we sampled were estimated to share a common ancestor less than 25 million years ago (Siler et al., 2012). Given that the pairs within each genus are closely related and have similar life histories, we do not expect substantive differences in mutation rates. Nonetheless, small differences in rates would affect the comparability of our divergence time estimates across our comparisons. Because we assumed a mutation rate of one for all comparisons, the estimated time of divergence for each pair of populations would still be accurate in units of expected substitutions per site. However, the assumption that the relative estimates among pairs are proportional to absolute time would be violated. Thus, at least some of the variation in divergence times we estimated among taxa is due to variation in rates of mutation.

A strong assumption about relative rates of mutation must be made for any comparative phylogeographic method to compare the timing of events across taxa (Hickerson et al., 2006; Huang et al., 2011; Chan et al., 2014; Oaks, 2014, 2018). This is because there is no information in the data to distinguish differences in mutation rates among comparisons when the population history of each is modeled separately (i.e., they are modeled as disconnected “species” trees). To relax this assumption, fully phylogenetic approaches to the problem of estimating co-divergences are needed so that information from the data about relative mutation rates across the phylogeny can inform the model while jointly estimating co-divergences.

#### Assumptions about migration

We also assumed there was no migration between the populations of each pair after they diverged. One reason for this assumption is practical: Currenty, `ecoevolity` does not model migration. Methods for estimating shared divergences that allow migration are based on approximate likelihoods and cannot handle genomic data (Huang et al., 2011; Oaks, 2014). Even when there is no migration, these methods have been shown to be extremely sensitive to prior assumptions and biased toward estimating shared divergences (Oaks et al., 2013; Hickerson et al., 2014; Oaks et al., 2014; Oaks, 2014). A primary cause of this poor behavior is that the insufficient summary statistics used by these methods contain little information about the divergence times and population sizes. Adding additional migration parameters to these models is likely to make inference more challenging; i.e., trying to estimate additional parameters with insufficient statistics. Although ignoring migration is not ideal, it allows us to use genomic data with a full-likelihood method that exhibits much more desirable statistical behavior than approximate alternatives.

The second reason for assuming no migration is biological; given the insularity and natural history of these geckos, we do not expect contemporary migration between pairs of islands to be an important process. Undoubtedly, these geckos have dispersed among islands of the Philippines, but such over-water dispersal events are likely too rare to meaningfully contribute to contemporary gene flow. Nonetheless, gene flow among connected islands during glacial periods certainly could have been significant. A biologically inspired model of migration would thus require assumptions about (1) periods of time when island exposure was conducive for migration, (2) divergences pre-dating these periods to allow migration during them, and (3) *absolute* rates of mutation to ensure the molecular evolution of both genera is on the same timescale as the divergence and subsequent periods of potential migration. As a result, modeling the timing and magnitude of repeated bouts of migration would be challenging and would necessarily ignore a lot of uncertainty, especially regarding absolute mutation rates. Also, the data would likely lack information about processes back beyond the most recent bout of migration. Instead, we can more simply estimate the last time each pair of populations experienced significant gene flow (i.e., the divergence time modeled by `ecoevolity`). This is a simpler inference problem, and the climate-driven PAIC model of vicariant divergence would still predict clustering among the most recent time the population pairs diverged. Furthermore, it seems unlikely that migration during glacial periods would bias our approach toward recovering independent divergences.

#### Assumptions about population sizes

Although we allowed the ancestral and descendant populations to have different effective sizes, we assumed these sizes were constant through time. This assumption was likely violated, especially if many of the divergences between the pairs of populations we sampled were due to over-water dispersal. Such events would likely have a strong founder effect on the effective size of one of the descendant populations. These demographic changes could affect our estimates of divergence times, however, based on coalescent theory, there are two reasons these effects should not be very large. First, the effective size of (the rate of coalescence within) the ancestral population is the most influential on the divergence time, because it determines the lag between the population divergence and the final gene coalescences, the latter of which is what the genetic data directly inform. Thus, as long as we have a reasonable estimate of the effective size of the ancestral population before the divergence, we should be able to accurately estimate the time of divergence. Second, the error in divergence-time estimation caused by demography should be limited to a magnitude on the order of ≈ 2*N*_*e*_ (or 2*N*_*e*_*μ* in the current study), because this is the expected disparity between gene coalescence and population divergence. Therefore, the additional variation in the data that is explained by incorporating more demographic realism may be offset by the error introduced by the additional model complexity.

### Performance of ecoevolity with empirical RADseq data?

Given the caveats discussed above, and those associated with any model-based inference, it is important to evaluate how well the method implemented in `ecoevolity` can estimate divergence times conditional on the RADseq data we sampled from the gekkonid populations. Perhaps, with so many genomic data, `ecoevolity` fails to accurately estimate uncertainty in divergence times, and thus is biased toward finding differences between comparisons that do not exist. However, when we included two comparisons that represented random subsets of the loci from the same pair of populations in the analysis, `ecoevolity` strongly supported that they co-diverged. Thus, given real RADseq data from two comparisons that co-diverged, `ecoevolity` can confidently place them together.

Furthermore, from analyzing 2,000 data sets that were simulated to match the dimensions of our gekkonid RADseq data, we found that `ecoevolity` was able to accurately and precisely estimate the timing (Figure 6) and number (Figure 7) of divergence events. Also, these results confirm the findings of Oaks (2018) that the method performs better when analyzing all sites rather than only unlinked SNPs. This is important, because it shows this behavior generalizes to data sets simulated to match the linkage and missing data patterns of empirical RADseq data.

It is not surprising that `ecoevolity` performs better when using all of the data, despite the linkage among sites within loci violating the model. This behavior matches theoretical expectations that the parameters in the model should not be biased by the linked sites, because information about linkage among sites is not used by the model. Linkage among sites does not change the expected site patterns under the model, it only reduces the variance of those patterns. Thus, the accuracy of estimates of divergence times and effective sizes of populations should not be affected by linked sites, as demonstrated here and by Oaks (2018). Furthermore, removing all but (at most) one variable site per locus is a rather draconian measure to avoid violating the linkage assumption, because it discards a substantial amount of informative data. Our results are also consistent with those of Chifman and Kubatko (2014), who found quartet inference of species trees from SNP data was also robust to the violation of unlinked characters. Nonetheless, our results should not be generalized to other methods that assume unlinked characters, especially methods that use information about site linkage patterns.

Our simulation results also show that `ecoevolity` is robust to large disparities in the number of sampled individuals and loci (Table 1; Figures 6 & 7). This is not surprising given Oaks (2018) found the benefit of collecting more characters begins to plateau quickly, even with as few as 200 loci. For example, we can compare the estimation of divergence times between the pairs that were simulated to match our RADseq data from *C. philippinicus* populations from the islands of Luzon and Babuyan Claro versus Polillo and Luzon. The former pair consists of only two samples per population and 3,855 loci, whereas the latter has five samples per population and 19,561 loci (Table 1). Despite these large disparities in sampling, the accuracy and precision of divergence-time estimates are very similar, especially when all sites are included in the analyses (Figure S17).

Perhaps most importantly, our simulation results also allow us to better interpret our empirical findings. There are two patterns worth highlighting in this regard. (1) When applied to datasets that were simulated with all eight pairs diverging independently (the rightmost column of the plots in Figure 7), `ecoevolity` has only moderate success in preferring a model with eight divergence events. (2) When the true number of divergences is less than eight, `ecoevolity` almost never estimates eight divergences (only 3 out of almost 2,000 simulations; Figure 7). Taken together, these observations demonstrate that `ecoevolity` is unlikely to spuriously support the model where all pairs of populations diverge independently. Thus, the empirical support we found for all eight pairs of *Cyrtodactylus* and *Gekko* populations diverging independently is likely robust.

### Sensitivity to the prior on divergence times

It is interesting that in analyses of both genera we see support for shared divergences increase as the prior on divergence times becomes more diffuse (Figures S5 & S11). Although less extreme here, this is the same pattern seen in approximate-likelihood Bayesian approaches to this problem (Oaks et al., 2013; Hickerson et al., 2014; Oaks et al., 2014). Hickerson et al. (2014) proposed this pattern was caused by numerical problems, whereas Oaks et al. (2014) interpreted the problem as being more fundamental: as more prior density is placed in regions of divergence-time space where the likelihood tends to be low, models that have fewer divergence-time parameters have greater marginal likelihoods because their likelihoods are “averaged” over less space with low likelihood and substantial prior weight. Our results clearly support the latter explanation, as the MCMC approach used here does not suffer from the insufficient prior sampling proposed by Hickerson et al. (2014).

Whereas the full-likelihood Bayesian approach used here is much more robust to prior assumptions than the ABC approaches, our results demonstrate that it is still important to assess sensitivity of the results to the priors (Oaks et al., 2013). This is especially true for the posterior probabilities of divergence models or the number of divergence events, which are the result of the prior probabilities being updated by the marginal likelihoods of the divergence models. Because the marginal likelihoods are averaged with respect to the priors on all the parameters of the model, they can be sensitive to those priors regardless of the informativeness the data (Oaks et al., 2019).

### Gekkonid diversification in the Philippines

Philippine *Gekko* and *Cyrtodactylus* species are nocturnal insectivores that inhabit a variety of geological substrates, forest types, and variable local atmospheric conditions (e.g., prevailing temperatures and precipitation) throughout many Philippine landmasses where they are codistributed. The Philippine *Gekko* populations studied here exhibit a more restricted microhabitat preference for rocky substrates, and appear more patchily distributed in the vicinity of exposed rock, caves, and karst formations. *Cyrtodactylus* are also found in some of these same habitats, but also utilize forest interior microhabitats, where they perch additionally on tree trunks and understory vegetation. Until recently, widespread species were recognized in both genera (e.g., *Cyrtodactylus philippinicus* and *Gekko mindorensis*), suggesting their microhabitat preferences do not limit their vagility. However, subsequent investigations of widespread taxa have shown Philippine gekkonid species diversity to be underestimated greatly and represented by a larger number of range-restricted lineages (Brown et al., 2009, 2011; Linkem et al., 2010b; Welton et al., 2010a,b).

Our findings are consistent with what we know about Philippine gekkonid natural history (RMB and CDS pers. obs.) and on-going revisions of their species boundaries. The spatial and temporal variation in connectivity among pockets of these lizards’ preferred structural microbabitats is likely a key predictor of past and present disributions of gekkonid populations across the Philippines. Environmental heterogeneity within islands is likely important for isolating populations, as evidenced by previous findings of multiple divergent lineages inhabiting the same island, such as the northern island of Luzon (Siler et al., 2010, 2012, 2014; Welton et al., 2010a,b). Furthermore, we consider it likely that the ephemeral, low-elevation habitat on the land bridges exposed during glacial periods was unsuitable for these forest species (Esselstyn and Brown, 2009; Hosner et al., 2014). In fact, rare, long-distance dispersal events among islands might actually be more likely to occur via rafting on vegetation across marine barriers following typhoons than movement across exposed land bridges during glacial periods (Linkem et al., 2013; Brown, 2016). Both intra-island processes of isolation associated with spatial and temporal environmental heterogeneity (Brown et al., 2013) and inter-island rafting (Brown, 2016) would predict our results of idiosyncratic divergence times across inter-island pairs of populations.

### Conclusions

Climate-driven fragmentation of the Philippine Islands has been invoked as a model of pulsed co-speciation throughout the archipelago. This model predicts that population divergences between fragmented islands should be temporally clustered around interglacial rises in sea levels. We analyzed comparative genomic data from 16 pairs of insular gecko populations within a full-likelihood, Bayesian model-choice framework to test for shared divergence events. Our results support independent divergences among the pairs of gecko populations. Although comparative genomic data from more taxa will allow us to address additional questions, our results suggest the repeated cycles of climate-driven island fragmentation has not been an important shared mechanism of speciation for gekkonid lizards in the Philippines.

## Supporting information

Animation for Supplemental Figure 2

## Data Accessibility

All of the demultiplexed, raw sequence reads are available from the NCBI Sequence Read Archive (Bioproject PRJNA486413, SRA Study SRP158258). A detailed history of all aspects of this project was recorded in a version-controlled repository, which is publicly available at https://github.com/phyletica/gekgo.

## Acknowledgments

We thank the members of the Phyletica Lab (the phyleticians) for helpful feedback on multiple drafts of this paper. We also thank editors Mohamed Noor and David Weisrock and four anonymous reviewers for constructive feedback that greatly improved this work. We are grateful to Patrick Monnahan and John Kelly for their help with the MSG library protocol. The computational work was made possible by the Auburn University (AU) Hopper Cluster supported by the AU Office of Information Technology and a grant of high-performance computing resources and technical support from the Alabama Supercomputer Authority. This work was possible thanks to funding provided to JRO from the National Science Foundation (DBI 1308885 and DEB 1656004). Philippines research has been supported by NSF DEB 0073199, 0743491, 0640737, 1418895, and 1654388 to RMB; NSF IOS 1353683 and DEB 1657648 and 0804115 to CDS; and the University of Kansas Biodiversity Institute. This paper is contribution number 891 of the Auburn University Museum of Natural History.

## Author contributions

J.R.O., C.D.S., and R.M.B. conceived the study. R.M.B. and C.D.S. conducted fieldwork. J.R.O. collected, assembled, and analyzed the sequence data. J.R.O. led the writing of the manuscript, with contributions from C.D.S. and R.M.B.

## Supporting Information

Table S1. The data for all samples included in the three RADseq libraries are included in a separate tab-delimited text file (https://raw.githubusercontent.com/phyletica/gekgo/master/tex/tables/msg-samples-edited.txt).

Table S2. The data for all samples included in the 16 pairs of populations analyzed in this study are included in a separate tab-delimited text file (https://raw.githubusercontent.com/phyletica/gekgo/master/tex/manuscripts/ecoevolity/tables/comparison-samples-edited.txt). This is a subset of the data in Table S1.

**Table S3.**
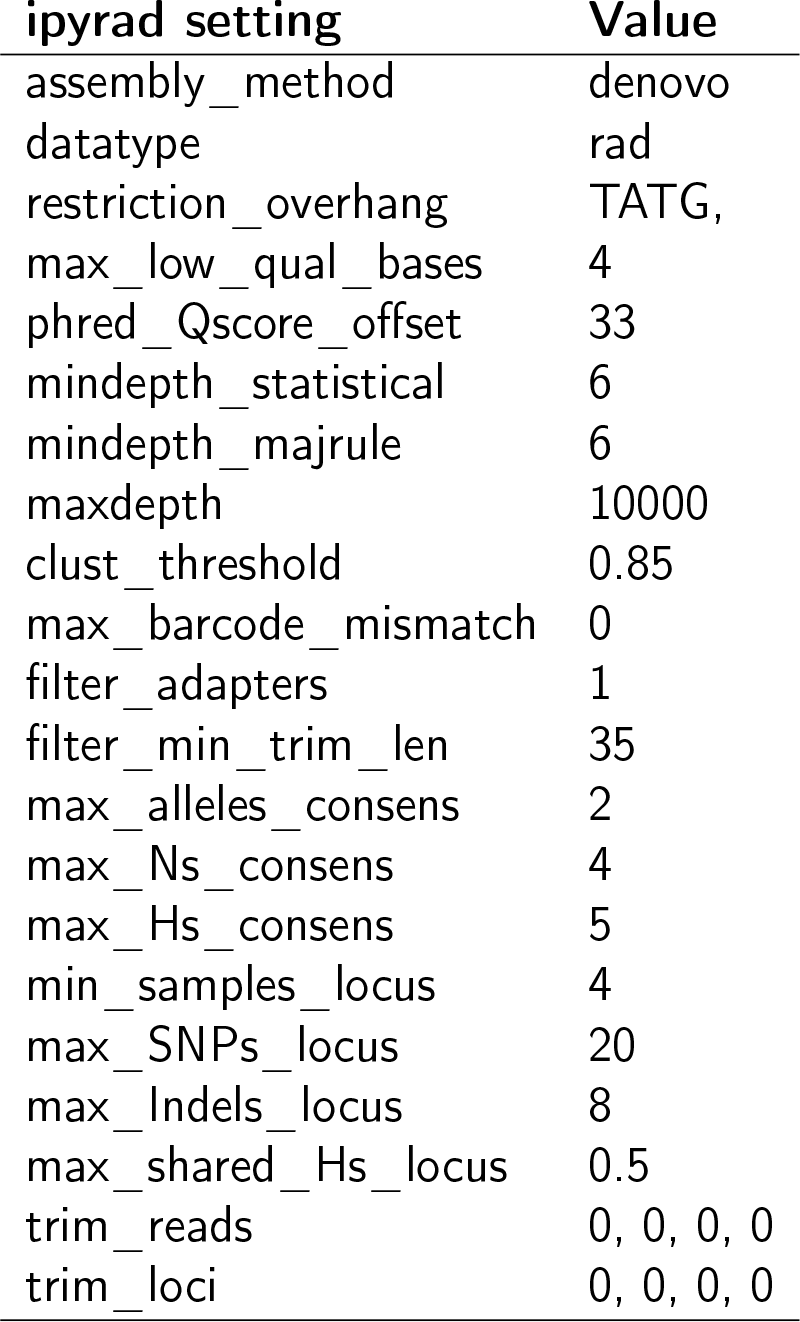
Settings used for assembling loci for each pair of gekkonid populations.

| **ipyrad setting** | **Value** |
| --- | --- |
| assembly_method | denovo |
| datatype | rad |
| restriction_overhang | TATG, |
| max_low_qual_bases | 4 |
| phred_Qscore_offset | 33 |
| mindepth_statistical | 6 |
| mindepth_majrule | 6 |
| maxdepth | 10000 |
| clust_threshold | 0.85 |
| max_barcode_mismatch | 0 |
| filter_adapters | 1 |
| filter_min_trim_len | 35 |
| max_alleles_consens | 2 |
| max_Ns_consens | 4 |
| max_Hs_consens | 5 |
| min_samples_locus | 4 |
| max_SNPs_locus | 20 |
| max_Indels_locus | 8 |
| max_shared_Hs_locus | 0.5 |
| trim_reads | 0, 0, 0, 0 |
| trim_loci | 0, 0, 0, 0 |

**Table S4.** Per-site nucleotide diversity within (*π*_1_ and *π*_2_) and between (*π*_*between*_) pairs of *Cyrtodactylus* and *Gekko* populations, calculated from the RADseq data using the SeqSift Python package (https://github.com/joaks1/SeqSift), which relies on Biopython (https://biopython.org/). The within-population nucleotide diversity (*π*_1_ and *π*_2_) is followed by the number of individuals sampled from the population in parentheses.

| Species | Island 1 | Island 2 | $\pi_1$ | $\pi_2$ | $\pi_{between}$ |
| --- | --- | --- | --- | --- | --- |
| *C. annulatus* | Bohol | Camiguin Sur | 0.00118 (4) | 0.00136 (4) | 0.00303 |
| *C. baluensis-redimiculus* | Palawan | Kinabalu | 0.00135 (4) | 0.00432 (3) | 0.02475 |
| *C. gubaot-sumuroi* | Samar | Leyte | 0.00573 (5) | 0.00223 (5) | 0.00653 |
| *C. philippinicus* | Luzon | Babuyan Claro | 0.00131 (2) | 0.00028 (2) | 0.00226 |
| *C. philippinicus* | Luzon | Camiguin Norte | 0.00102 (3) | 0.00068 (4) | 0.00349 |
| *C. philippinicus* | Polillo | Luzon | 0.00226 (5) | 0.00211 (5) | 0.00429 |
| *C. philippinicus* | Panay | Negros | 0.00177 (3) | 0.00095 (2) | 0.00335 |
| *C. philippinicus* | Sibuyan | Tablas | 0.00136 (3) | 0.00209 (3) | 0.00319 |
| *G. crombota-rossi* | Babuyan Claro | Calayan | 0.00101 (5) | 0.00111 (5) | 0.00340 |
| *G. gigante* | South Gigante | North Gigante | 0.00167 (3) | 0.00127 (4) | 0.00149 |
| *G. mindorensis* | Lubang | Luzon | 0.00183 (5) | 0.00119 (4) | 0.00851 |
| *G. mindorensis* | Panay | Masbate | 0.00137 (4) | 0.00130 (3) | 0.00407 |
| *G. mindorensis* | Negros | Panay | 0.00103 (3) | 0.00177 (5) | 0.00259 |
| *G. porosus* | Sabtang | Batan | 0.00092 (4) | 0.00115 (4) | 0.00172 |
| *G. romblon* | Romblon | Tablas | 0.00178 (5) | 0.00218 (2) | 0.00349 |
| *G. sp. a-sp. b* | Camiguin Norte | Dalupiri | 0.00096 (5) | 0.00111 (5) | 0.00262 |

**Figure S1.**
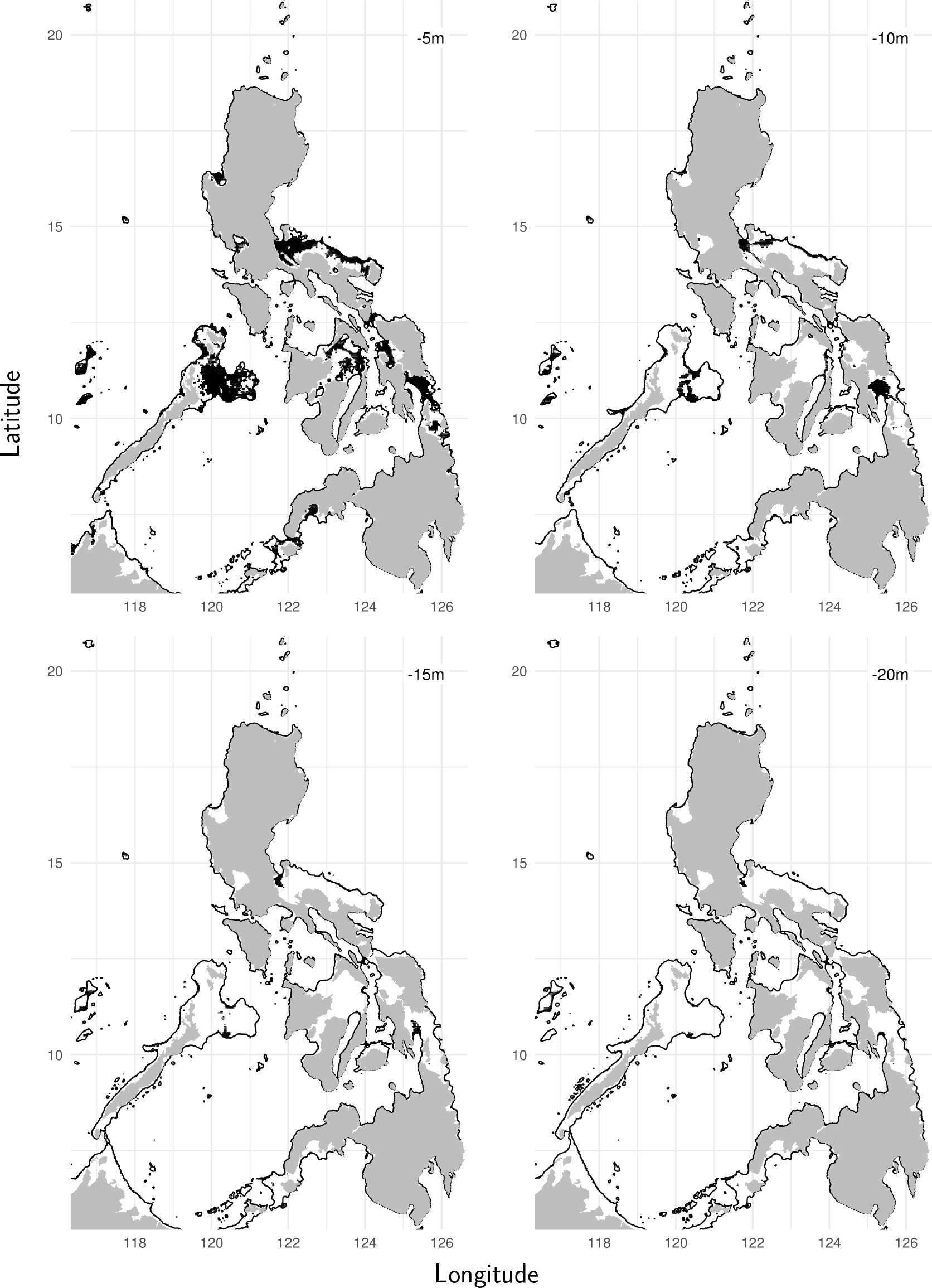
Bathymetry contours around the Philippine Islands at varying depths. Black lines show the contour associated with the depth indicated in the upper right of each plot. Contours are based on data from the ETOPO1 1-arc-minute global relief model (Amante and Eakins, 2009). Figure generated with marmap version 1.0.2 (Pante and Simon-Bouhet, 2013) and ggplot2 Version 2.2.1 (Wickham, 2009).

**Figure S2.**
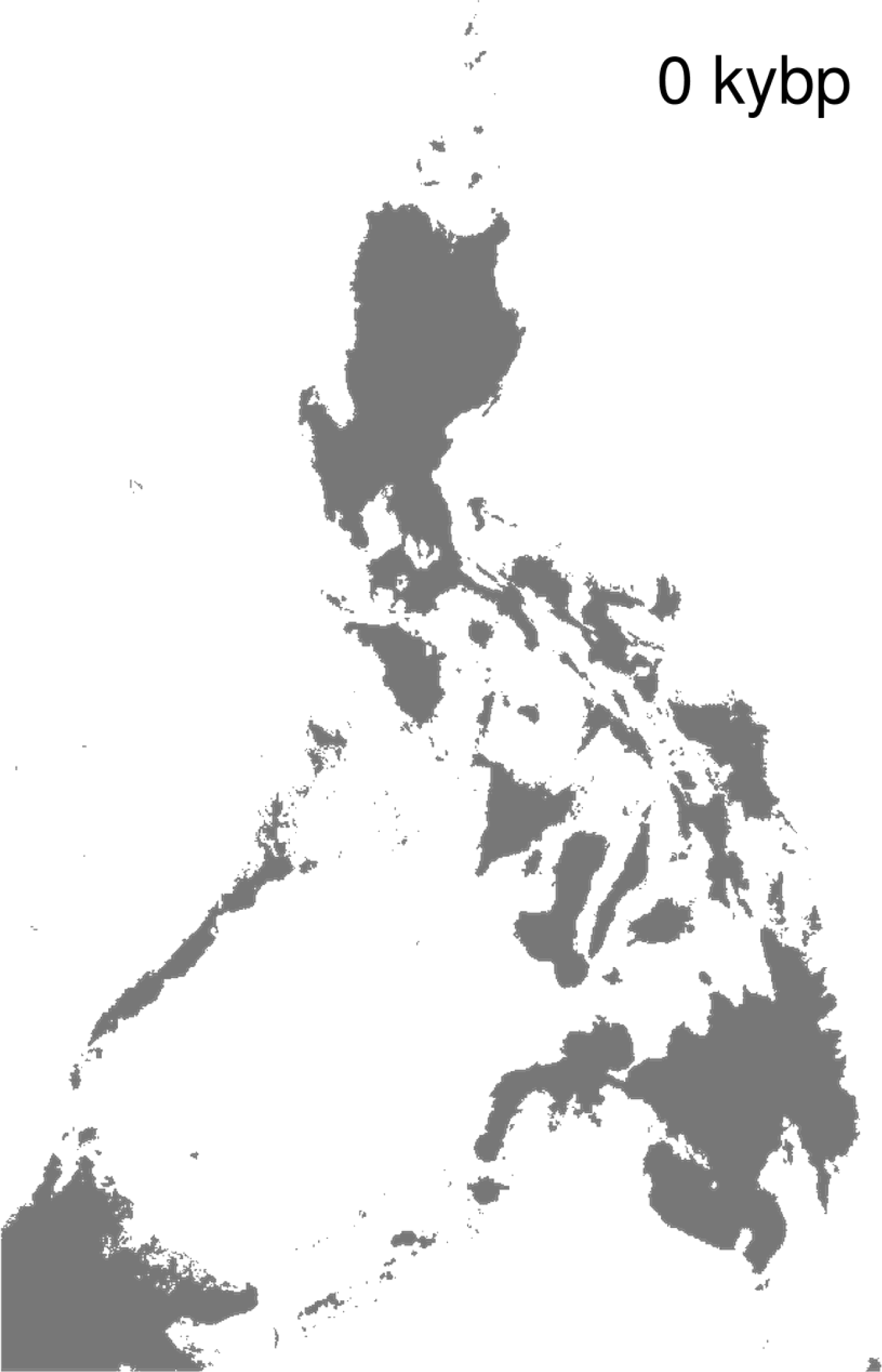
Animation of approximate sea-level changes in the Philippine Islands over the last 430,000 years. Sea-level estimates are from the projection of Spratt and Lisiecki (2016) based on seven reconstructions. Bathymetry data are from the ETOPO1 1-arc-minute global relief model (Amante and Eakins, 2009). Animation generated using marmap version 1.0.2 (Pante and Simon-Bouhet, 2013), ggplot2 Version 2.2.1 (Wickham, 2009), ImageMagick Version 6.9.10-8 Q16 x86_64 20180723, and FFmpeg Version 4.0.2-2. The source code for generating the plot is available at https://github.com/phyletica/animating-sea-level-change. This animation can also be viewed at https://youtu.be/NjGdCezUvw8.

**Figure S3.**
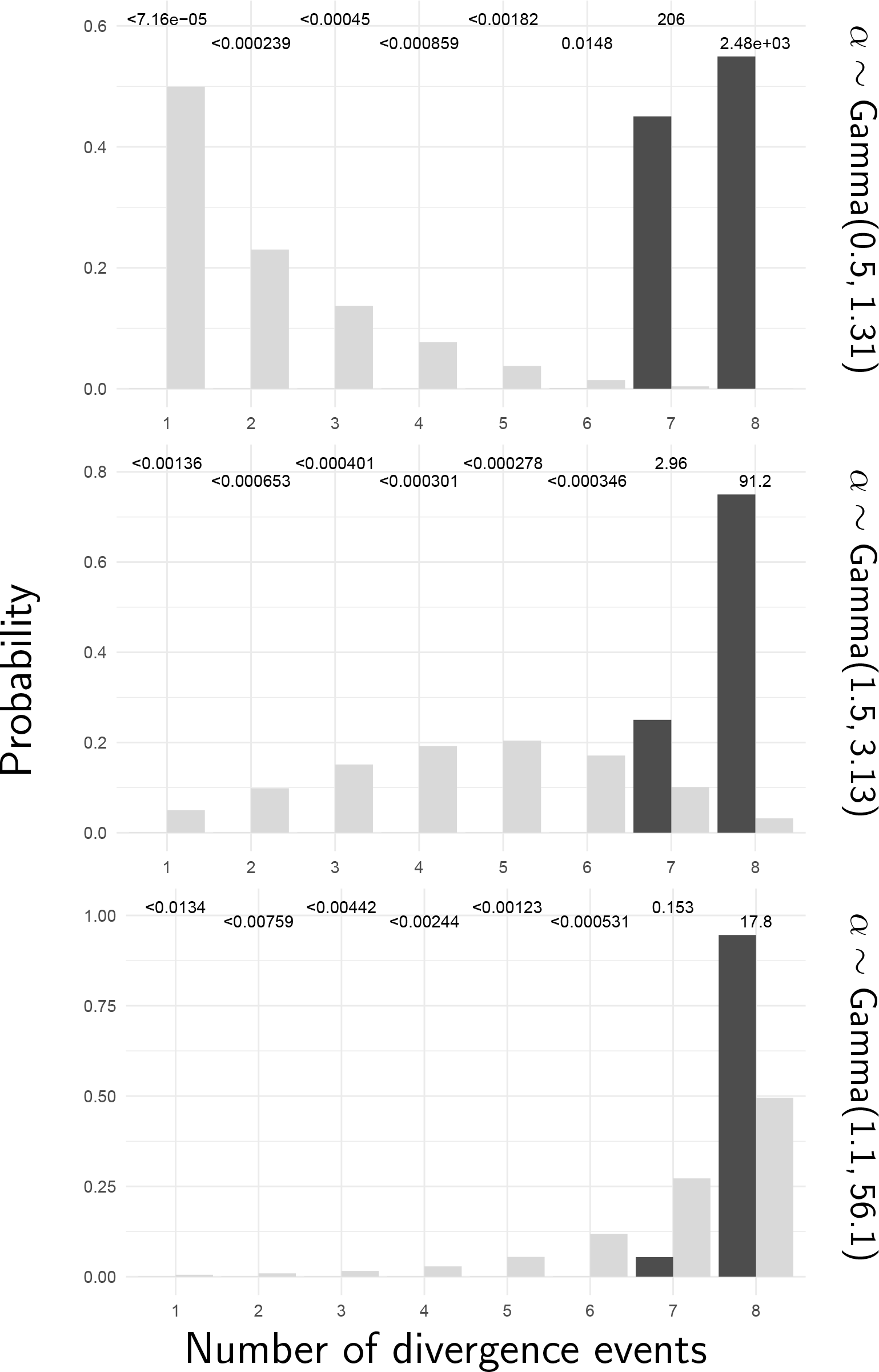
Approximate prior (light bars) and posterior (dark bars) probabilities of numbers of divergence events across pairs of *Cyrtodactylus* populations under three different priors on the concentration parameter of the Dirichlet process. Bayes factors for each number of divergence times is given above the corresponding bars. Each Bayes factor compares the corresponding number of events to all other possible numbers of divergence events. Figure generated with ggplot2 Version 2.2.1 (Wickham, 2009).

**Figure S4.**
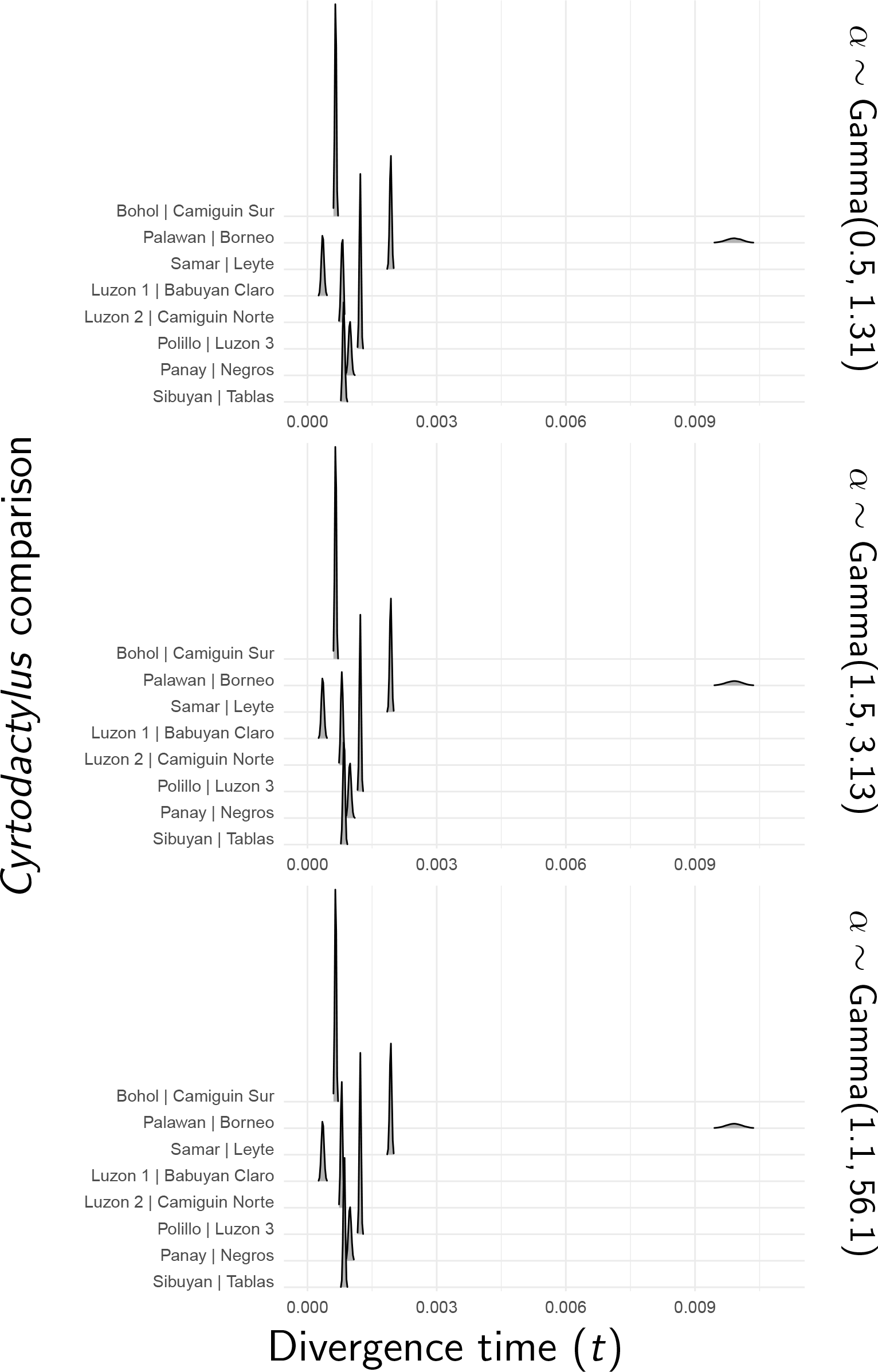
Approximate marginal posterior densities of divergence times for each pair of *Cyrtodactylus* populations under three different priors on the concentration parameter of the Dirichlet process. Figure generated with ggridges Version 0.4.1 (Wilke, 2018) and ggplot2 Version 2.2.1 (Wickham, 2009).

**Figure S5.**
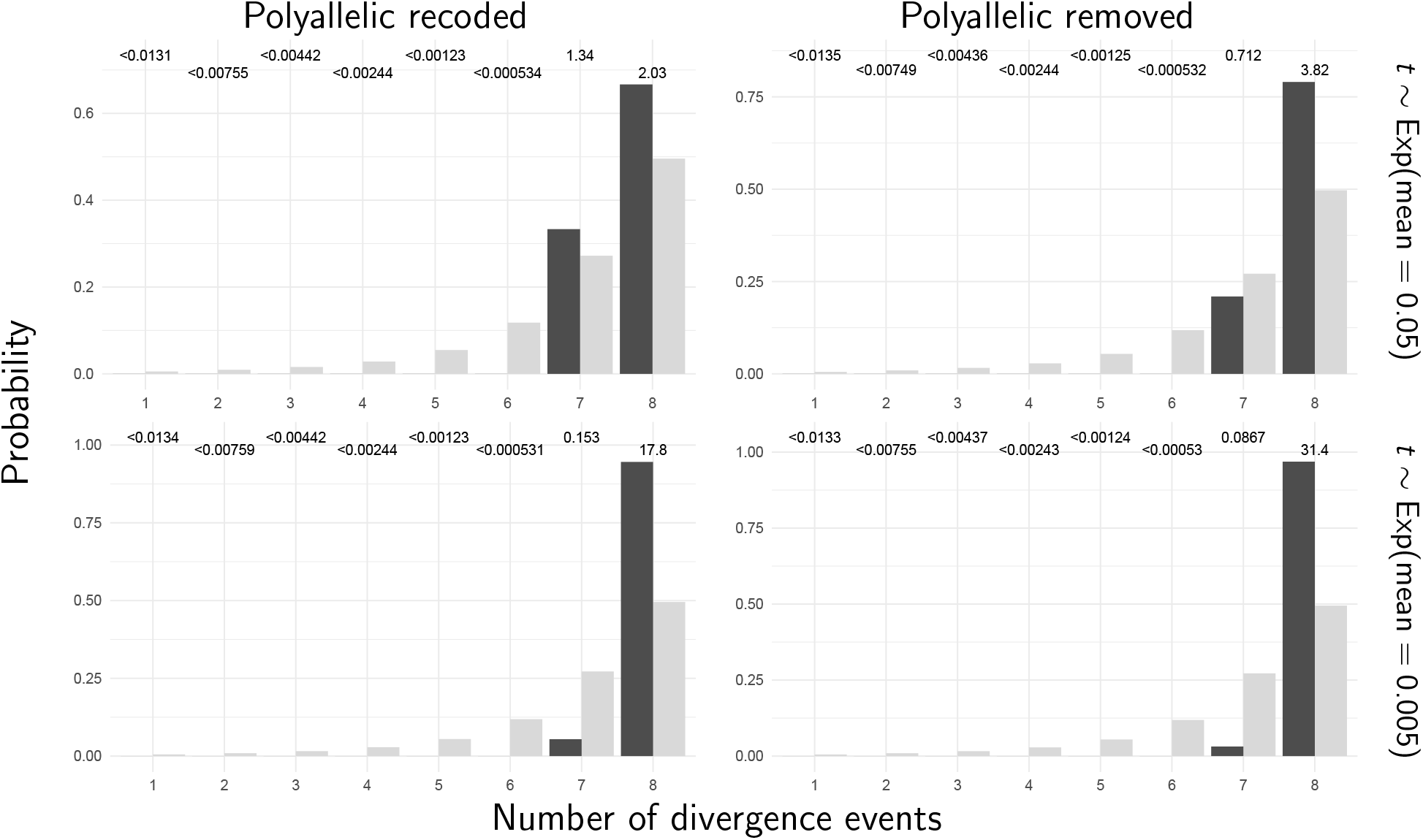
Approximate prior (light bars) and posterior (dark bars) probabilities of numbers of divergence events across pairs of *Cyrtodactylus* populations under four different combinations of prior on divergence times (rows) and recoding or removing polyallelic characters (columns). Bayes factors for each number of divergence times is given above the corresponding bars. Each Bayes factor compares the corresponding number of events to all other possible numbers of divergence events. Figure generated with ggplot2 Version 2.2.1 (Wickham, 2009).

**Figure S6.**
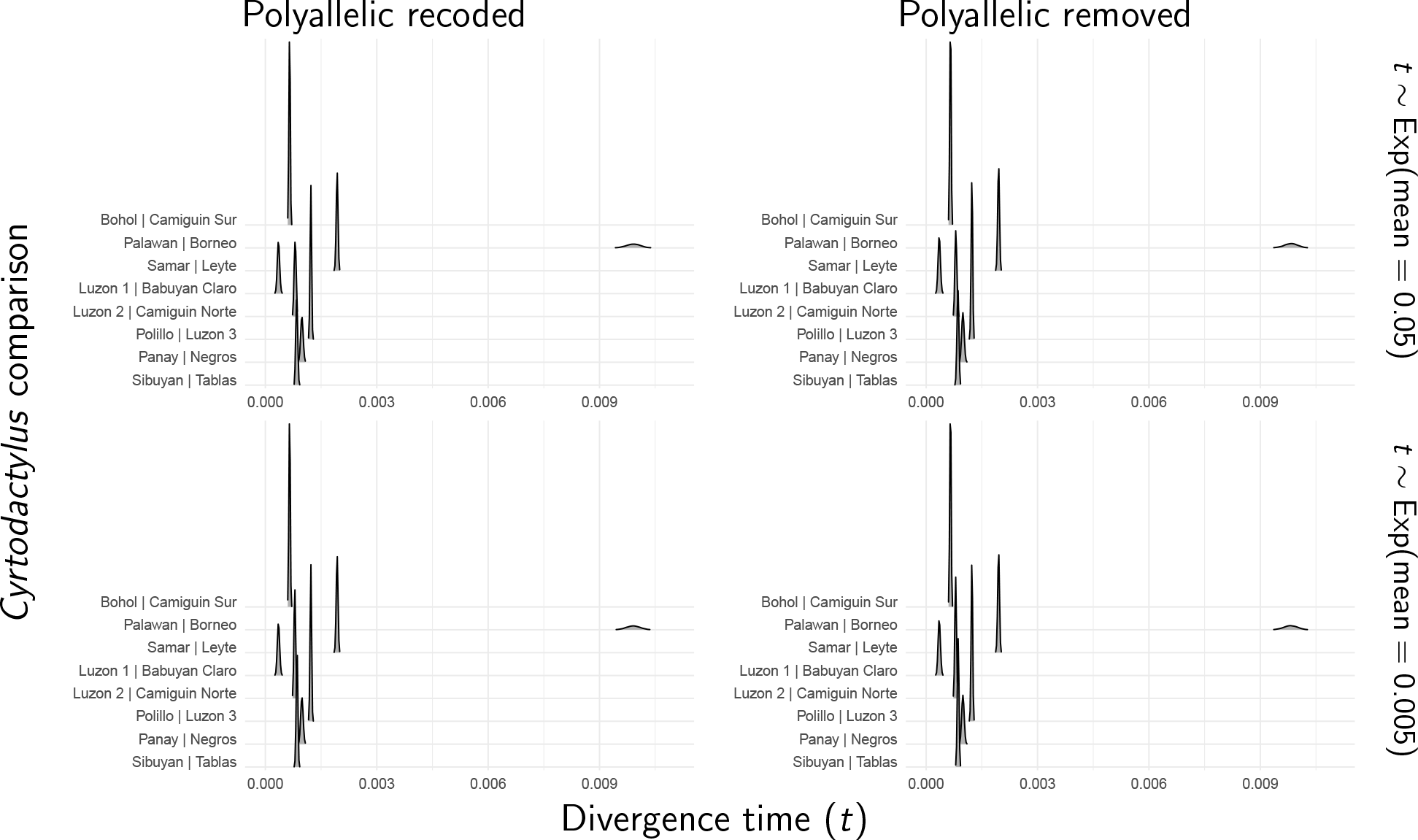
Approximate marginal posterior densities of divergence times for each pair of *Cyrtodactylus* populations under four different combinations of prior on divergence times (rows) and recoding or removing polyallelic characters (columns). Figure generated with ggridges Version 0.4.1 (Wilke, 2018) and ggplot2 Version 2.2.1 (Wickham, 2009).

**Figure S7.**
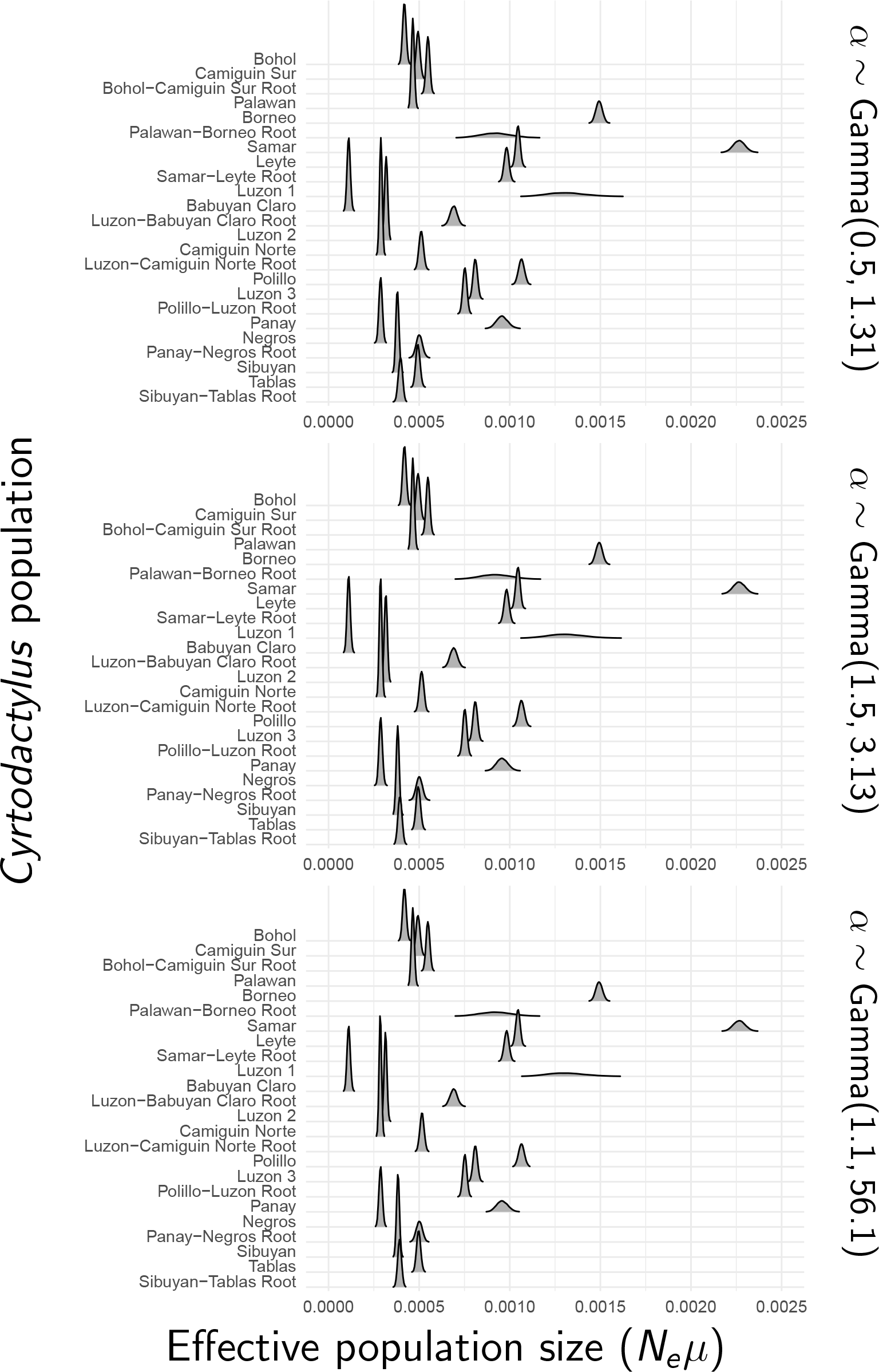
Approximate marginal posterior densities of population sizes for each pair of *Cyrtodactylus* populations under three different priors on the concentration parameter of the Dirichlet process. Figure generated with ggridges Version 0.4.1 (Wilke, 2018) and ggplot2 Version 2.2.1 (Wickham, 2009).

**Figure S8.**
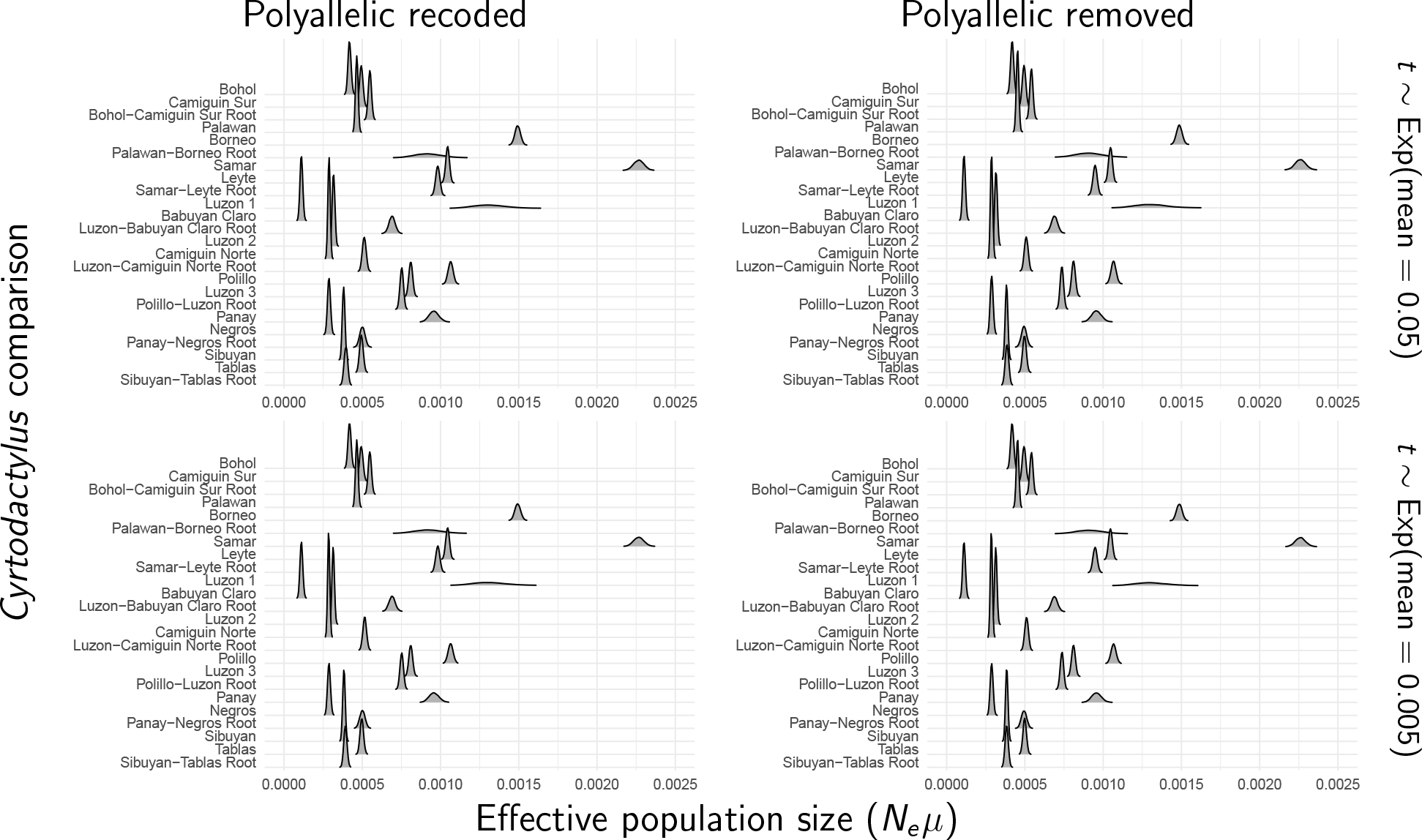
Approximate marginal posterior densities of population sizes for each pair of *Cyrtodactylus* populations under four different combinations of prior on divergence times (rows) and recoding or removing polyallelic characters (columns). Figure generated with ggridges Version 0.4.1 (Wilke, 2018) and ggplot2 Version 2.2.1 (Wickham, 2009).

**Figure S9.**
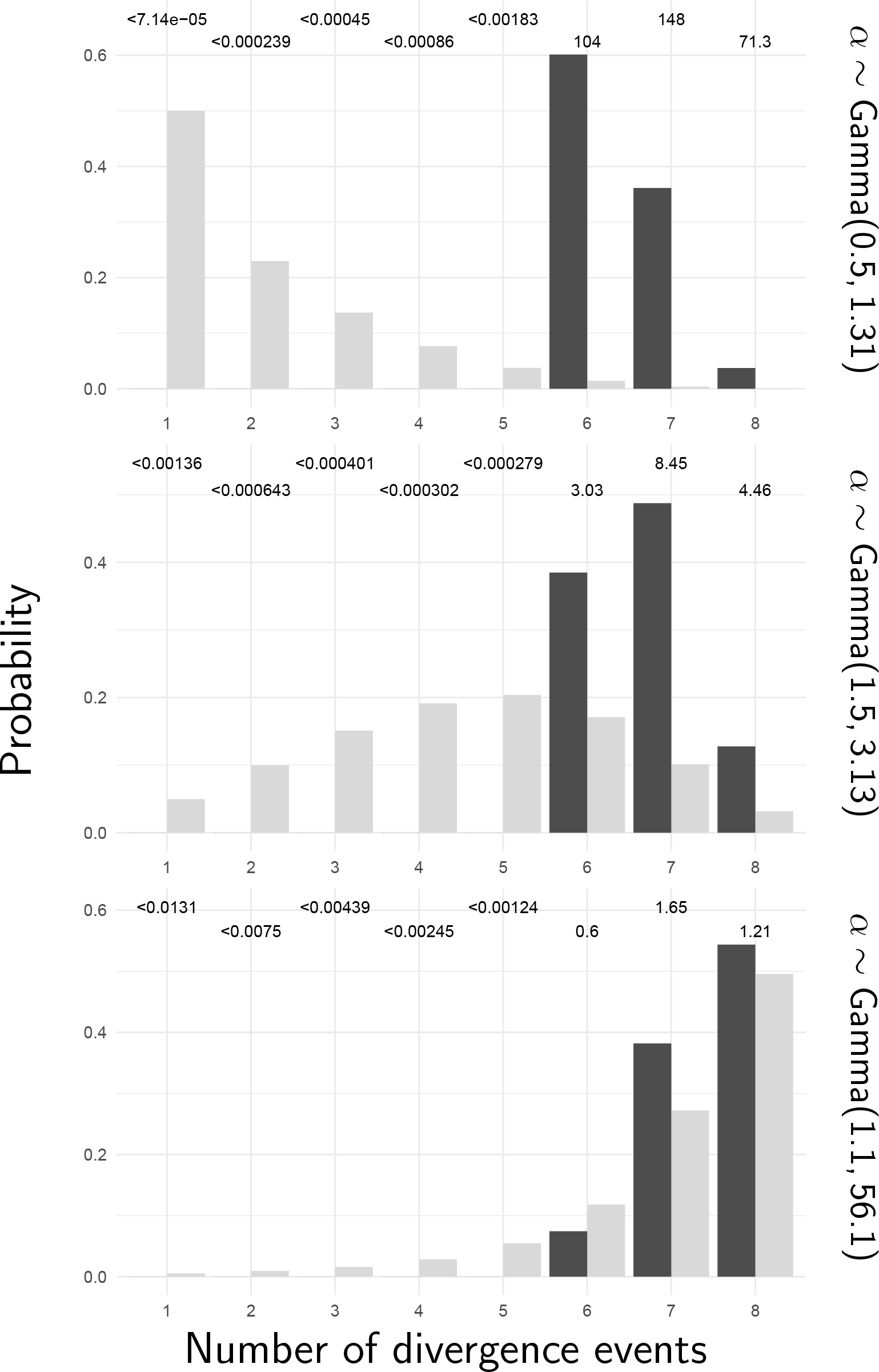
Approximate prior (light bars) and posterior (dark bars) probabilities of numbers of divergence events across pairs of *Gekko* populations under three different priors on the concentration parameter of the Dirichlet process. Bayes factors for each number of divergence times is given above the corresponding bars. Each Bayes factor compares the corresponding number of events to all other possible numbers of divergence events. Figure generated with ggplot2 Version 2.2.1 (Wickham, 2009).

**Figure S10.**
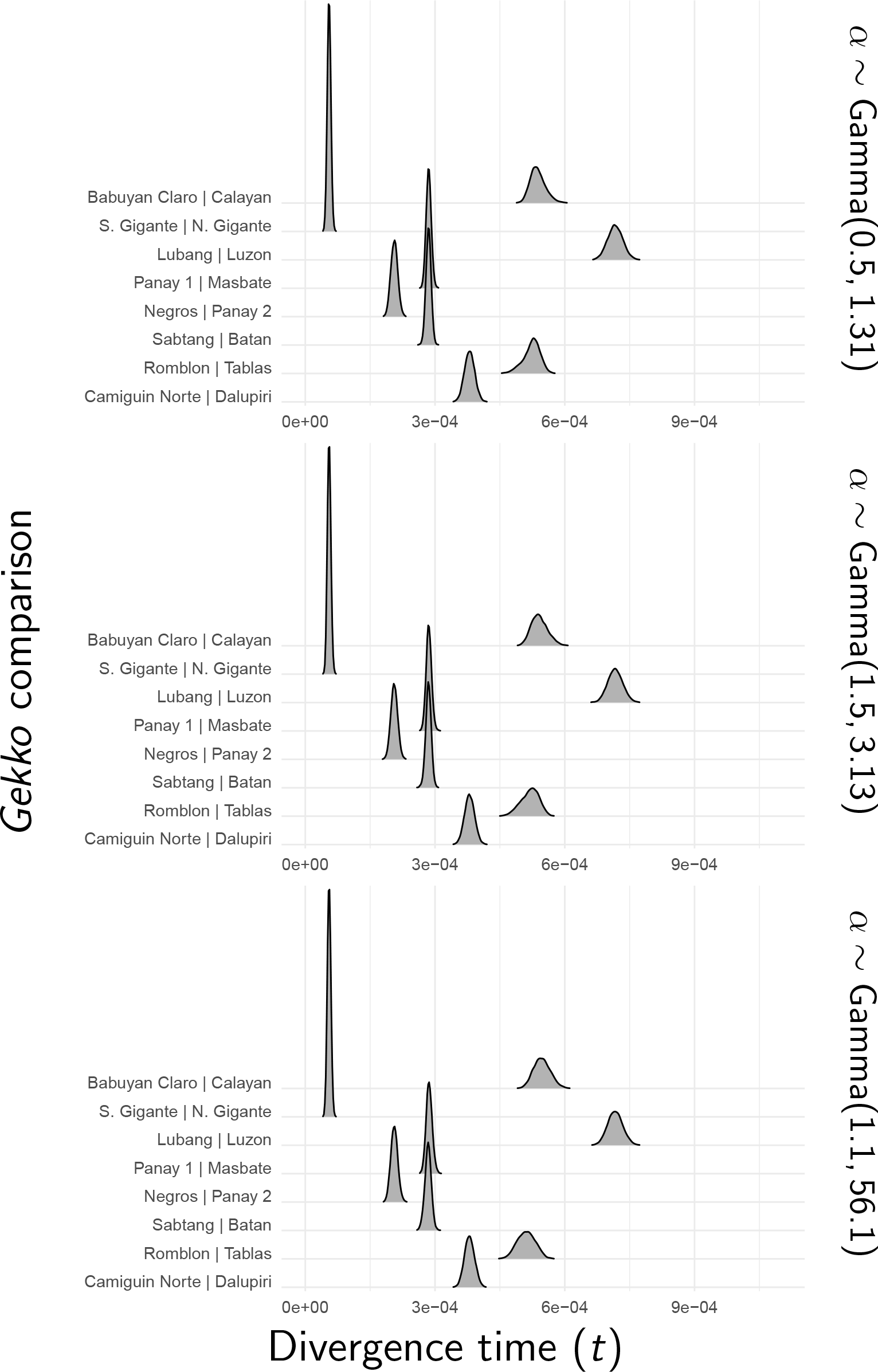
Approximate marginal posterior densities of divergence times for each pair of *Gekko* populations under three different priors on the concentration parameter of the Dirichlet process. Figure generated with ggridges Version 0.4.1 (Wilke, 2018) and ggplot2 Version 2.2.1 (Wickham, 2009).

**Figure S11.**
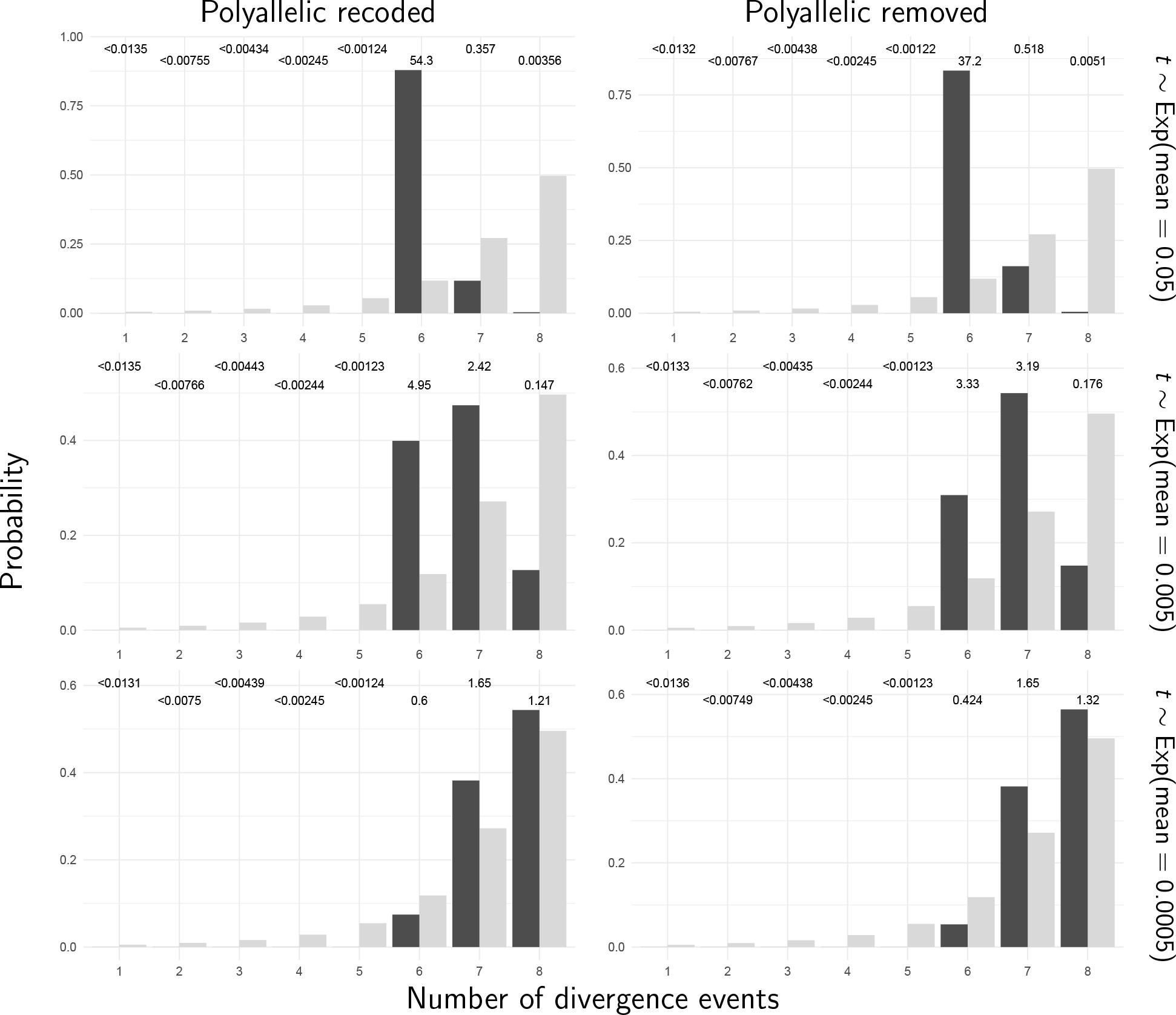
Approximate prior (light bars) and posterior (dark bars) probabilities of numbers of divergence events across pairs of *Gekko* populations under six different combinations of prior on divergence times (rows) and recoding or removing polyallelic characters (columns). Bayes factors for each number of divergence times is given above the corresponding bars. Each Bayes factor compares the corresponding number of events to all other possible numbers of divergence events. Figure generated with ggplot2 Version 2.2.1 (Wickham, 2009).

**Figure S12.**
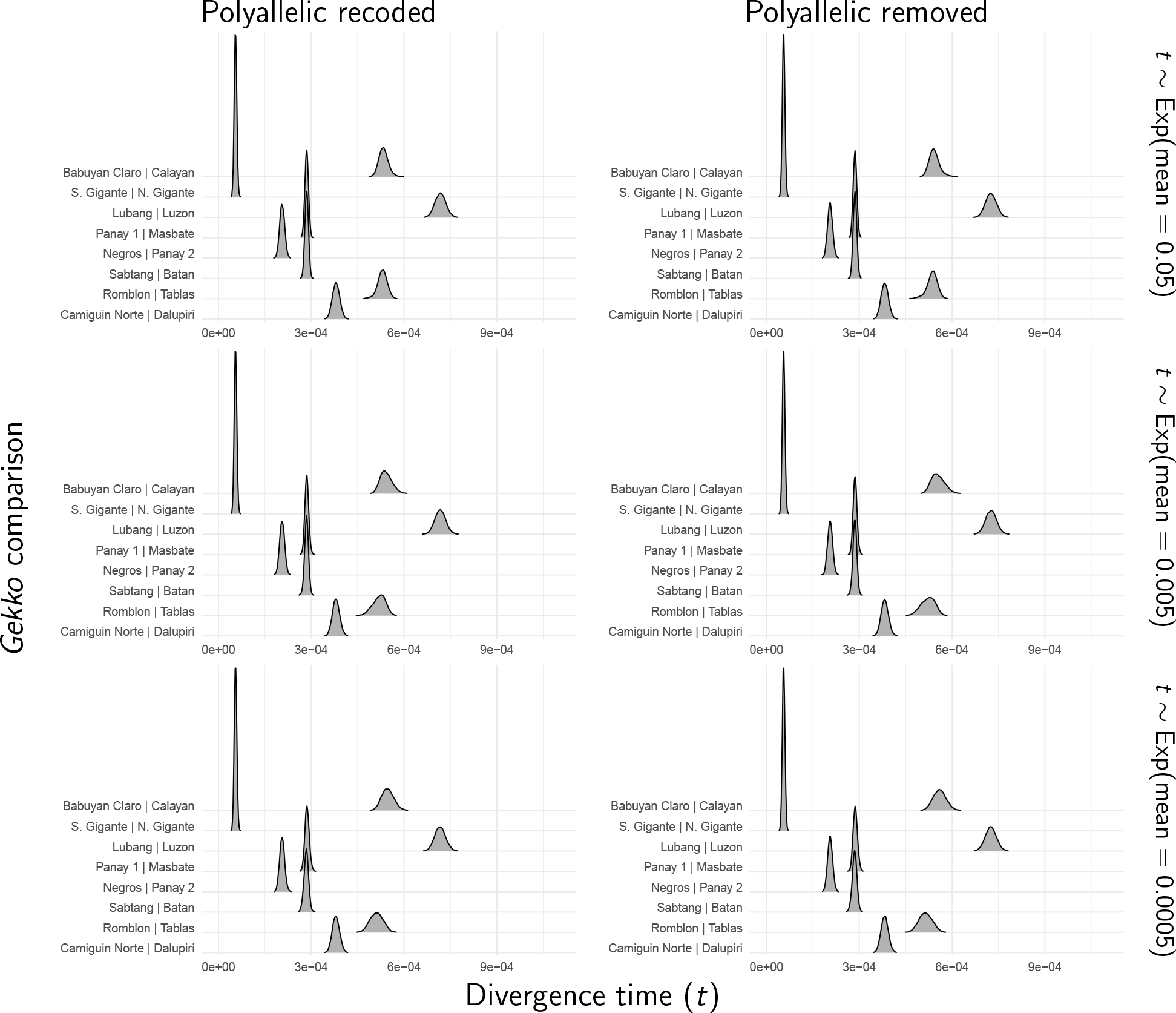
Approximate marginal posterior densities of divergence times for each pair of *Gekko* populations under six different combinations of prior on divergence times (rows) and recoding or removing polyallelic characters (columns). Figure generated with ggridges Version 0.4.1 (Wilke, 2018) and ggplot2 Version 2.2.1 (Wickham, 2009).

**Figure S13.**
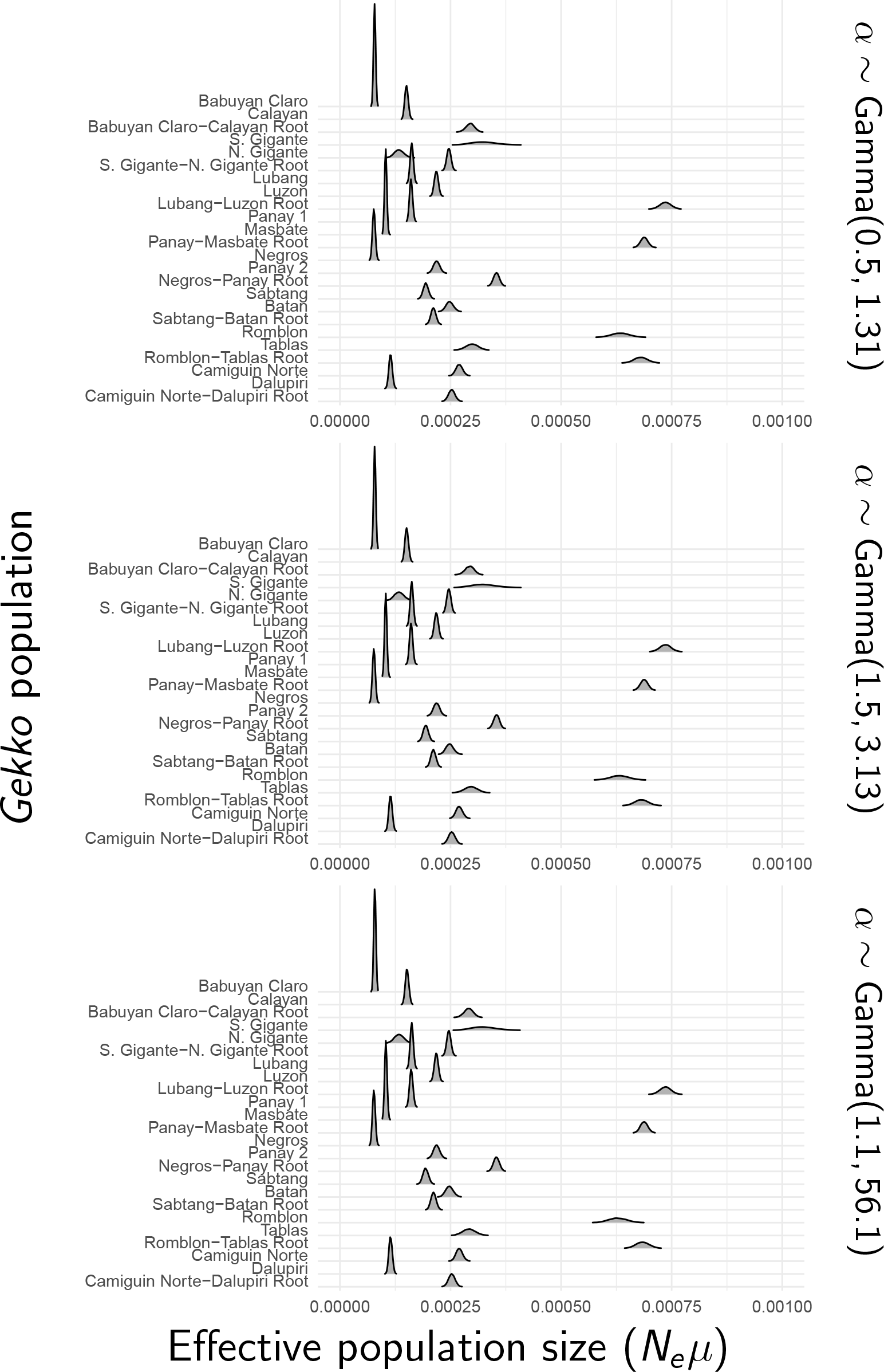
Approximate marginal posterior densities of population sizes for each pair of *Gekko* populations under three different priors on the concentration parameter of the Dirichlet process. Figure generated with ggridges Version 0.4.1 (Wilke, 2018) and ggplot2 Version 2.2.1 (Wickham, 2009).

**Figure S14.**
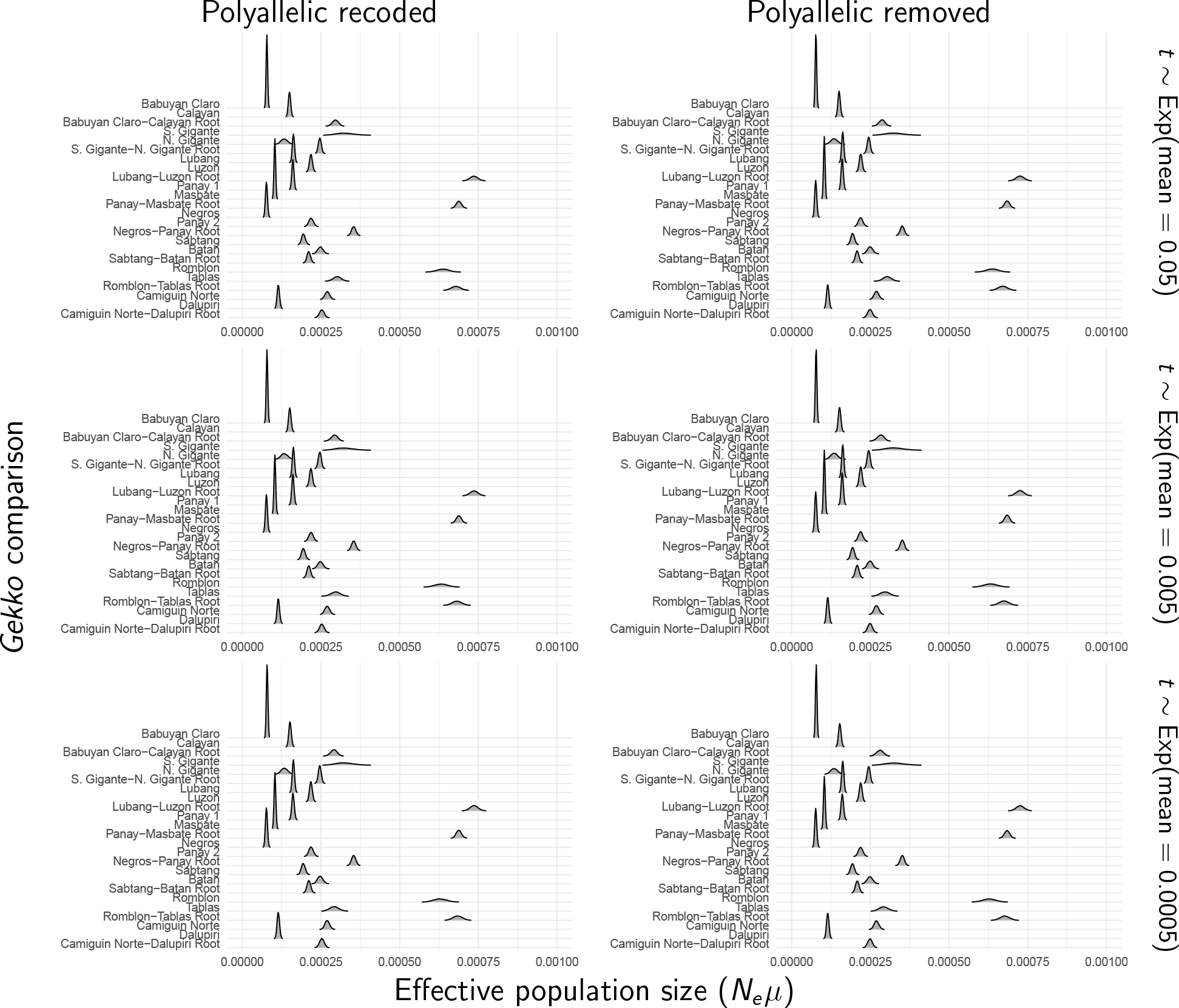
Approximate marginal posterior densities of population sizes for each pair of *Gekko* populations under six different combinations of prior on divergence times (rows) and recoding or removing polyallelic characters (columns). Figure generated with ggridges Version 0.4.1 (Wilke, 2018) and ggplot2 Version 2.2.1 (Wickham, 2009).

**Figure S15.**
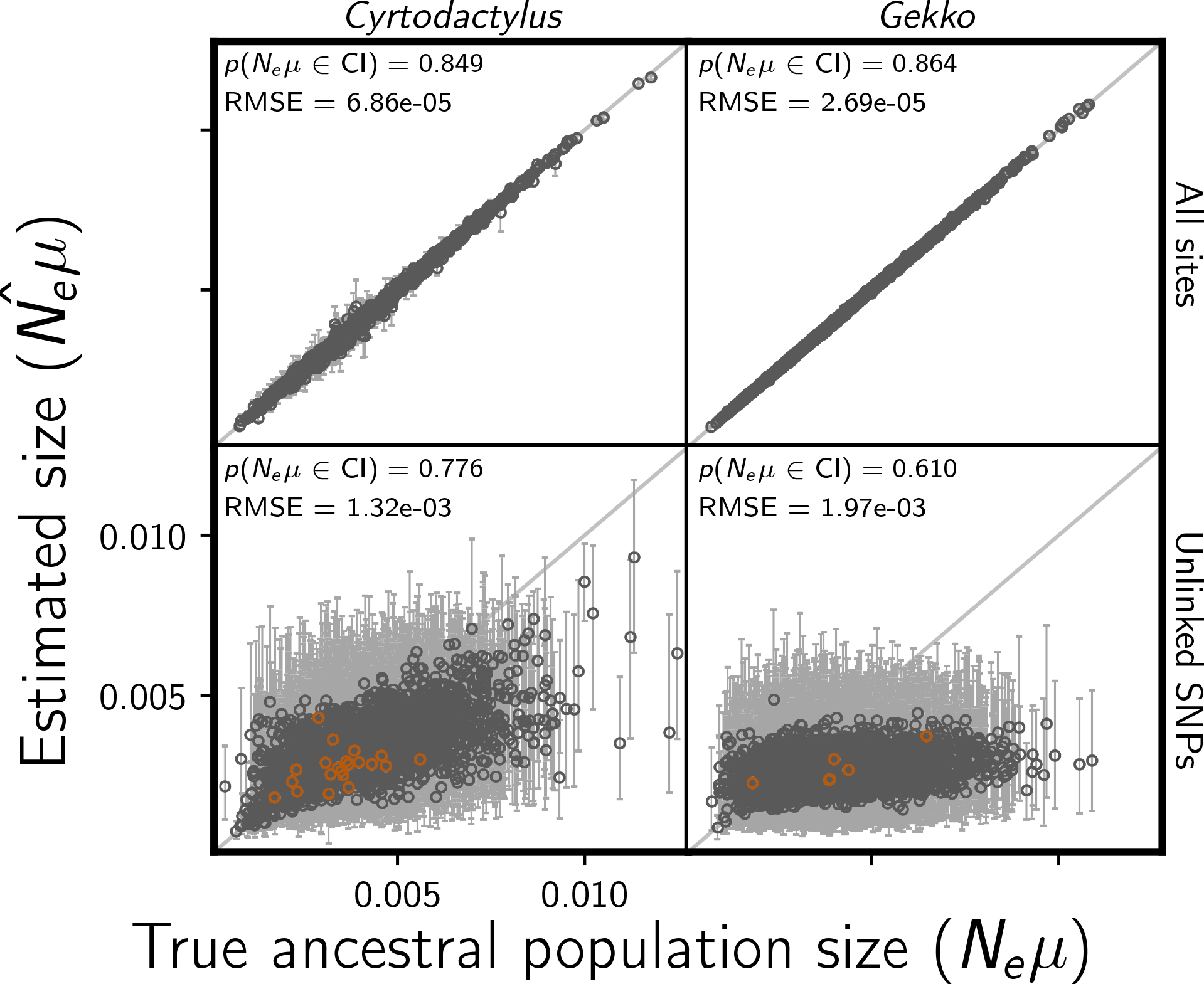
The accuracy and precision of `ecoevolity` estimates of the ancestral population size (scaled by the mutation rate) when applied to data simulated to match our *Cyrtodactylus* (left) and *Gekko* (right) RADseq data sets with all sites (top) or only one SNP per locus (bottom). Each circle and associated error bars represent the posterior mean and 95% credible interval. Estimates for which the potential-scale reduction factor was greater than 1.2 (Brooks and Gelman, 1998) are highlighted in orange. Each plot consists of 4000 estimates—500 simulated data sets, each with eight pairs of populations. For each plot, the root-mean-square error (RMSE) and the proportion of estimates for which the 95% credible interval contained the true value—*p*(*N*_*e*_*μ* ∈ CI)—is given. Figure generated with matplotlib Version 2.0.0 (Hunter, 2007).

**Figure S16.**
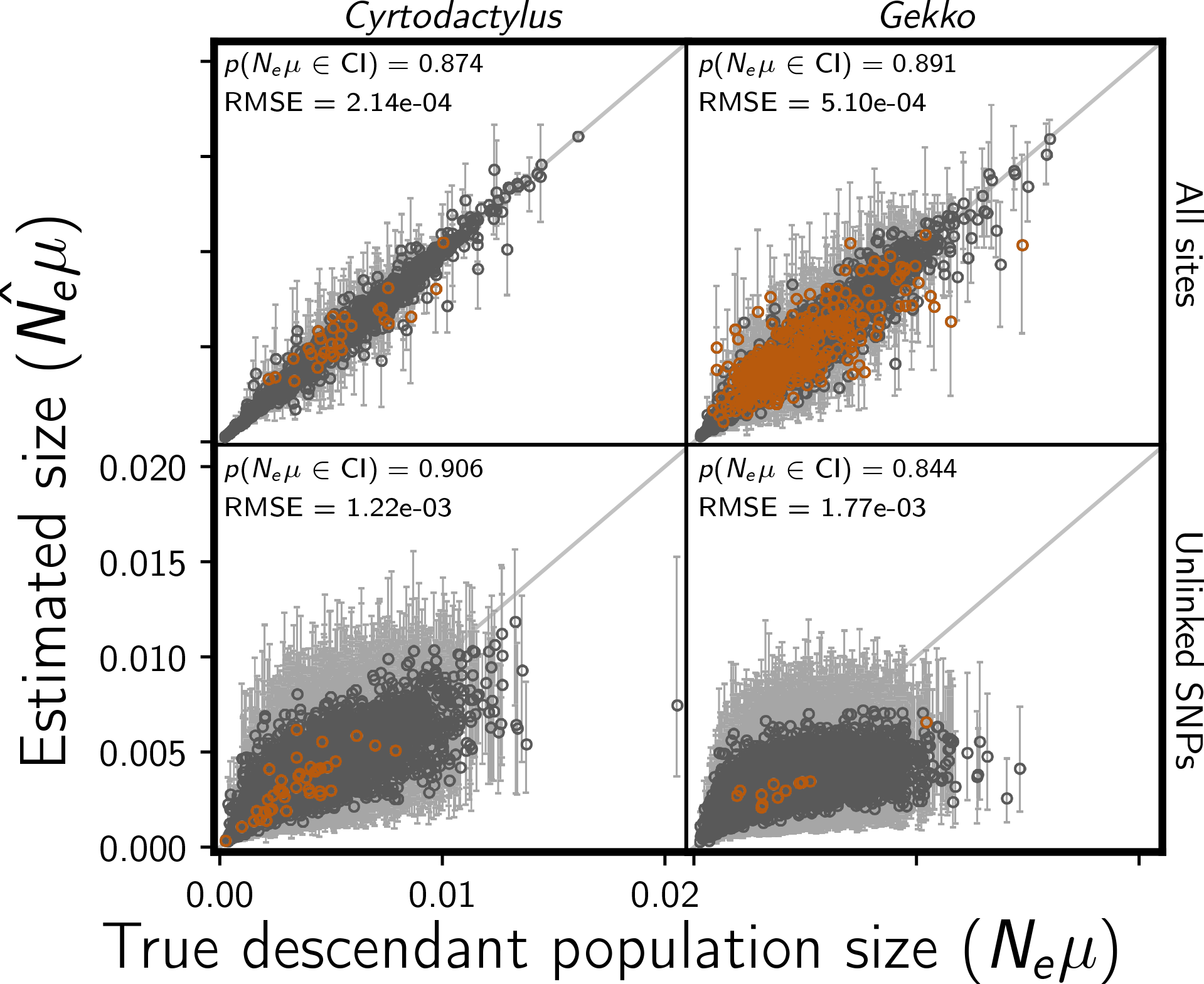
Accuracy and precision of `ecoevolity` estimates of the descendant population sizes (scaled by the mutation rate) when applied to data simulated to match empirical *Cyrtodactylus* (left) and *Gekko* (right) RADseq data sets with all sites (top) or only one SNP per locus (bottom). Each circle and associated error bars represents the posterior mean and 95% credible interval. Estimates for which the potential-scale reduction factor was greater than 1.2 (Brooks and Gelman, 1998) are highlighted in orange. Each plot consists of 8000 estimates—500 simulated data sets, each with eight pairs of populations. For each plot, the root-mean-square error (RMSE) and the proportion of estimates for which the 95% credible interval contained the true value— *p*(*N*_*e*_*μ* ∈ CI)—is given. Figure generated with matplotlib Version 2.0.0 (Hunter, 2007).

**Figure S17.**
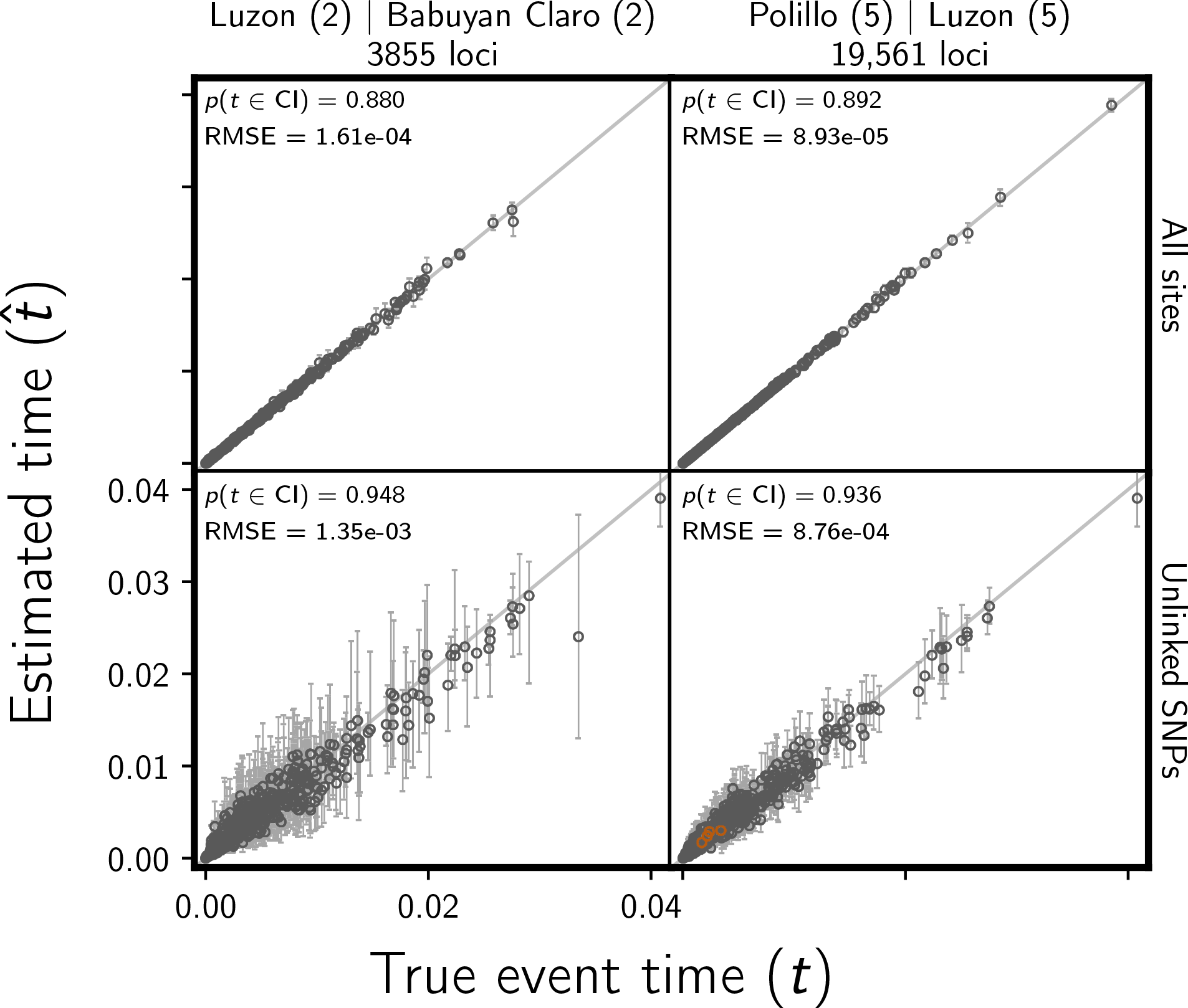
Accuracy and precision of `ecoevolity` divergence-time estimates (in units of expected subsitutions per site) when applied to data simulated to match empirical RADseq data sets sampled from the pairs of *Cyrtodactylus philippinicus* populations from the islands of (left) Luzon and Babuyan Claro and (right) Polillo and Luzon (a subset of the results plotted in Figure 6). Results are shown for ecoevolity analyses of data sets that contain all sites (top), or only one SNP per locus (bottom). The number of individuals sampled from each island population is indicated in parentheses at the top of each column of plots. Results for these two pairs of populations are plotted separately here to compare divergence-times estimated from data sets with large differences in the number of sampled individuals and loci. Each circle and associated error bars represents the posterior mean and 95% credible interval for the time that a pair of populations diverged. Estimates for which the potential-scale reduction factor was greater than 1.2 (Brooks and Gelman, 1998) are highlighted in orange. Each plot consists of 500 estimates—one from each of the 500 simulated data sets. For each plot, the root-mean-square error (RMSE) and the proportion of estimates for which the 95% credible interval contained the true value—*p*(*t* ∈ CI)—is given. Figure generated with matplotlib Version 2.0.0 (Hunter, 2007).

